# Selective vulnerability of intracortical projection neurons drives long-term cortical circuit rewiring after perinatal hypoxia

**DOI:** 10.64898/2026.09.21.753115

**Authors:** Salma Ellouze, Elodie Babina, Géraldine Meyer-Dilhet, Louis Foucault, Amel Amara, Stevenson Desmercière, Thomas Gagnieu, Sandrine Mongellaz, Anis Cataldi, Guillaume Marcy, Jean Francois Ghersi-Egea, Julien Courchet, Olivier Raineteau

**Affiliations:** Univ Lyon, Université Claude Bernard Lyon 1, Inserm, Stem Cell and Brain Research, Institute U1208, 69500 Bron, France; Univ Lyon, Univ Lyon 1, CNRS, INSERM, Physiopathologie et Génétique du Neurone et du Muscle, UMR5261, U1315, Institut NeuroMyoGène, 69008 Lyon, France; Univ Lyon, Université Claude Bernard Lyon 1, Labex CORTEX, bioinformatics platform, 69008 Lyon, France; Fluid Team, Lyon Neurosciences Research Center, INSERM U1028, CNRS UMR5292, Lyon University, 69500 Bron, France

## Abstract

Cortical circuits are built at perinatal times and gradually refined in an activity-dependent manner during a so-called postnatal period of critical plasticity. Although lesions of the central nervous system (CNS) happening during this period typically recover better than those occurring later in life, they are often associated with long-term behavioral deficits. This suggests that neuronal circuits rewiring, in particular within the cortex, may either be incomplete or inappropriate.

To address this possibility, we used chronic perinatal hypoxia, a mouse model of very premature birth. We confirmed that chronic hypoxia induced a decrease in cortical thickness frequently observed in very preterm babies, which rapidly recovered 8 days later. To explore the transcriptional correlates of this recovery we next performed single-nuclei transcriptomic analysis of the cortex at short (P11) and long (P45) timepoints following hypoxia. This revealed persistent transcriptional changes within neurons, including of genes involved in mitochondrial metabolism, axonogenesis and synaptogenesis. Further, histological analysis using anterograde and retrograde tracing as well as mitochondrial labelling support persistent alterations in upper cortical neurons resulting in increased corticocortical connectivity within the cortex of adult hypoxic mice. Finally, behavioral testing revealed altered social behavior in mice exposed to chronic hypoxia, which amplified with age.

Altogether, our results unravel how brain lesions happening early in life, alter normal cortical development and have long term consequences on cortical wiring, contributing to the observed behaviors defects appearing later in life.

## Introduction

The early postnatal period is a critical period of plasticity, when brain circuits are made, broken and refined under the influence of the organism’s environment ^1, 2^. It is also a period of intense neuritic growth and pruning, especially for layer 2/3 pyramidal neurons (PNs) which define cortico-cortical connectivity ^3^. The timing and stabilization of optimal cortical connectivity are governed by interacting mechanisms, including the maturation of GABAergic inhibition, particularly parvalbumin (PV) interneurons, which gate critical period onset and closure ^4, 5^. The progressive accumulation of extracellular matrix components, especially perineuronal nets, further restricts plasticity and consolidates synaptic configurations ^6^. Further, homeostatic scaling mechanisms ^7^ and thalamocortical dynamics ^8^ shape the balance between flexibility and stability during circuit refinement. Together, these mechanisms establish a temporally restricted but biologically coordinated framework that enables precise activity-dependent refinement of cortical connectivity while ensuring long-term circuit stability.

Premature babies have underdeveloped lungs that impair oxygenation, as well as periods of apnea due to immature respiratory control. These events can perturb cerebral oxygenation and systemic metabolic homeostasis, with potential short- and long-term consequences ^9^. Indeed, the early postnatal period relies on high metabolic demands that involve mitochondria ^10, 11^. This is because rapid brain growth, synaptogenesis, and circuit remodeling require substantial ATP produced primarily by mitochondrial oxidative phosphorylation ^10^. Importantly, mitochondrial and neuronal maturation are tightly interconnected and interdependent processes. As neurons differentiate and extend increasingly complex axonal and dendritic arbors, their mitochondria undergo profound changes in mass, morphology, dynamics, trafficking and oxidative capacity. In turn, these mitochondrial adaptations are required to sustain dendritic growth, synapse formation and maturation, by providing local energy and regulating calcium and redox homeostasis.^12–14^. Prematurity exacerbates vulnerabilities in mitochondrial maturation process, making energy dysregulation a core concern for early postnatal brain development and potential long-term outcomes.

Because prematurity disrupts cerebral maturation as well as the critical period of plasticity, it is often associated with abnormal circuits wiring as well as with the development of neuropsychiatric and behavioral deficits. Indeed, very premature babies are at increased risk of severe complications, including developmental delay, cerebral palsy, blindness or deafness, whose prevalence and severity are negatively correlated with gestational age and weight at birth (ref). Improved neonatal therapeutic care may alleviate some of the most severe clinical manifestations, yet leaving a higher predisposition to develop lasting neurodevelopmental impairment, including an increased risk for intellectual disabilities as well as autism spectrum disorders (ASD, 7 to 10X increase), bipolar disorders (7.4X) as well as other neurodevelopmental diseases ^15, 16^.

Here we used a chronic neonatal hypoxia murine model which recapitulates the physiopathological changes observed in premature, very low birth weight babies ^17^. By combining single-nucleus RNA sequencing with histological analyses, we identify upper-layer glutamatergic neurons, particularly layer 2/3 pyramidal neurons (PN) involved in intracortical connectivity, as a major and previously underappreciated target of earlylife hypoxic stress. Focusing on those neurons, we show 1)both acute and delayed effects of the hypoxic period on glutamatergic neuron mitochondria, 2) their maturation, 3) the development of altered cerebral connectivity and functions. Together, our results provide new mechanistic insights in the long-term consequences of metabolic stresses occurring early in life, on brain wiring.

## Results

### Analysis of hypoxia induced transcriptional changes within cortical neurons reveals sensitivity of upper cortical neurons

We set out to explore the long-term consequences of a metabolic insult resulting from chronic hypoxia, occurring during the critical period of plasticity. To this end, we used the chronic hypoxia model, in which newborn mice are exposed to reduced oxygen for a prolonged period of time (10% O2, from Postnatal day (P)3 to P11) ^17^. As described in previous studies, this model leads to diffuse brain lesions as well as an overall growth retardation^18–20^. We could indeed confirm a statistically significant decrease of body weight immediately at the end of the hypoxic period (−33%) (**Fig. 1A**), which remained significant at P19 but eventually returned to baseline by P45 (not shown). The reduced growth was paralleled at P11 by a reduction in brain volume (**Fig. 1A**), accompanied by a ventriculomegaly as well as a thinning of the cerebral cortical mantle (**Fig. 1B**). Interestingly, and as previously described ^21^, hypoxic mice compensated this delay of brain development, and by P19, cortical thickness was indistinguishable between the hypoxic and normoxic groups (**Fig. 1B**), thereby preceding full body growth recovery.

**Figure 1.**
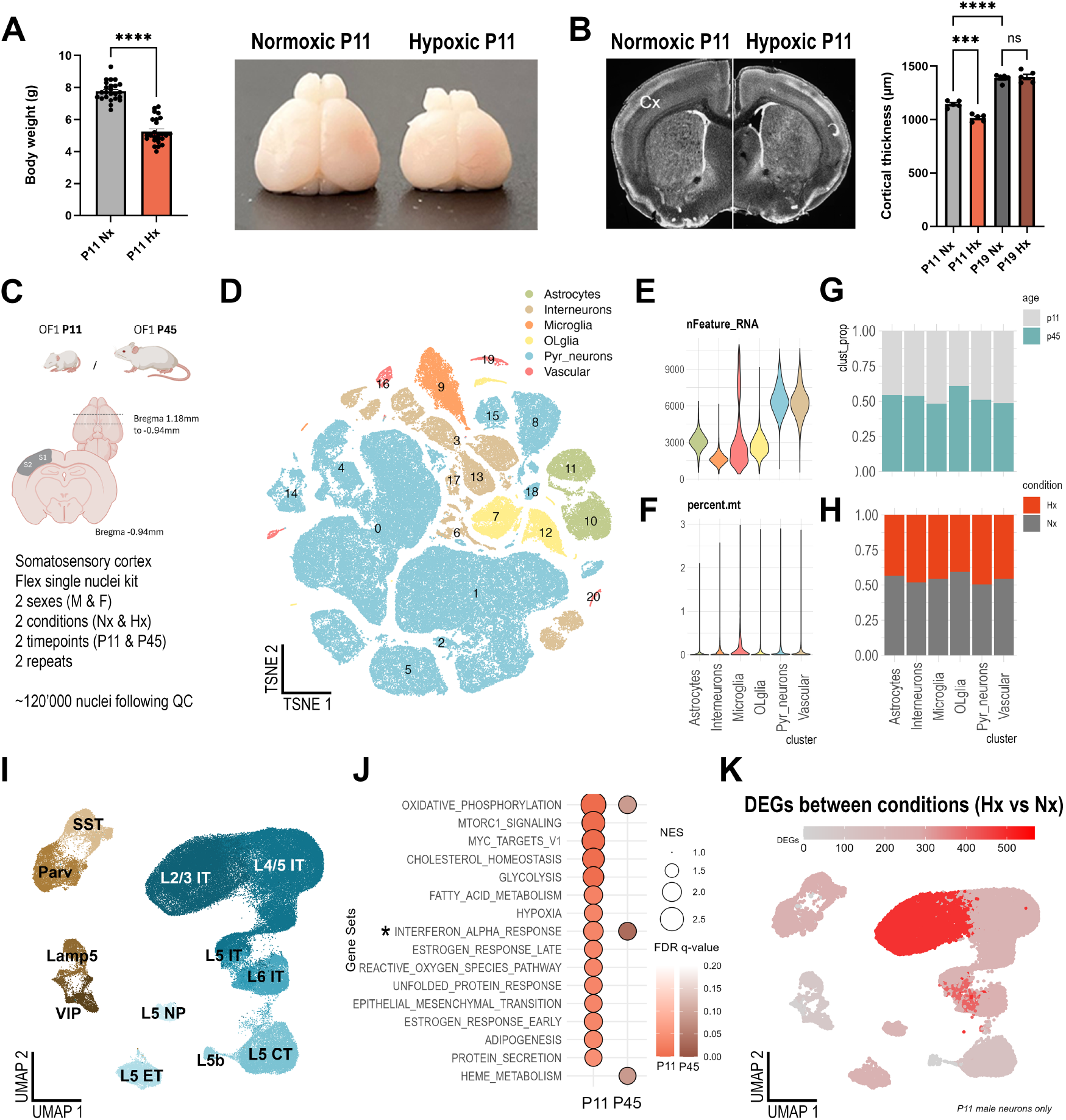
Large-scale single cell profiling identifies upper layers cortical neurons are primarily affected by postnatal hypoxia. **A)** chronic neonatal hypoxia impairs body weight at P11 as well as brain development. Statistical test: unpaired t test Mann-Whitney. Data: average±SEM. N_(P11Nx)_ = 24, N_(P11Hx)_ = 25. B) Enlargement of the ventricle as well as cortical thinning are observed on dapi counterstaining cross sections at P11 in hypoxic pups. Quantification reveals significant decrease of cortical thickness at P11, which recovers at P19. Large-scale single cell analysis of cortical cell types at P11 and P45 following hypoxia. Statistical test: two-way ANOVA with Bonferroni’s multiple comparison. Data: average±SEM. N_(P11Nx)_ = 5, N_(P11Hx)_ = 5, N_(P19Nx)_ = 5, N_(P19Hx)_ = 5. C) Scheme of the microdissection and characteristics of the multiplexing dataset. D) tSNE plot depicting the simplified identity annotation following QC. E-F) Summary of metrics of the dataset. Cells to be analyzed are selected based on the following criteria: nCount_RNA<125000 & percent.mt<3. G-H) Plots depicting the proportion of nuclei/cell type at P11 vs P45 (E) and from normoxic vs hypoxic mice. I) UMAP plot of subclustered neurons and predicted identity of interneurons and pyramidal neurons subtypes. J) Hierarchical network of HALLMARK genesets identified by GSEA analysis on all neurons at P11 and P45. Asterix indicate downregulation by hypoxia K) Projected DEGs by neuron subtypes reveal a higher effect of hypoxia onto superficial L2/3 intra-telencephalic pyramidal neurons. ns not significant, ***p ≤ 0.001, ****p≤ 0.0001.

To gain an in-depth overview of cortical cellular and molecular heterogeneity following hypoxia, we performed large-scale single-nucleus transcriptional profiling using the multiplexing Flex technology approach from the microdissected somatosensory cortex (**Fig. 1C** and **Fig. S1**). This region is particularly vulnerable to prematurity-related injury and hypoxic stress, both in humans and rodent models, and provides a well-defined framework to study cortical neuron diversity and maturation ^21–23^. We analyzed two developmental stages, i.e. P11 to capture the immediate effects of neonatal hypoxia, and P45 to assess long-term transcriptional consequences.

In total, our dataset represents 16 samples (2 timepoints x 2 sexes (M & F), 2 conditions (Hx and Nx) and x 2 independent repeats) (**Fig. 1C**), representing ~120’000 nuclei following QC (**Fig. S1**). Clustering analysis at low resolution (0.1) revealed 21 clusters among which 14 were neuronal and 5 corresponded to non-neuronal cells (i.e. Astrocytes, OPCs, Microglia, MOLs, Vascular cells, **Fig. 1D**). Neurons could be identified based on the expression of generic markers (**Fig. S2**) of GABAergic interneurons (Interneurons, *Gad1, Gad2, Erbb4*, 15’903 nuclei) or glutamatergic pyramidal neurons (*Glu* neurons, *Tbr1, Cux2, Satb2*, 88’427 nuclei). Quality control revealed averaged detected genes ranging from 1500 to 6000 gene/cell for Microglia and *Glu* neurons, respectively (**Fig. 1E**), with consistently low percentage of nuclear-encoded mitochondrial genes revealing successful isolation of nuclei (**Fig. 1F**). Proportion of nuclei were balanced between P11 and P45 for all cell types (**Fig. 1G**). Similarly, the proportion of nuclei isolated from normoxic and hypoxic mice were comparable for all cell type, allowing comparison of gene expression in both conditions (**Fig. 1H**).

Owing to the higher prevalence and severity of neurodevelopmental disorders in males following premature birth^24^, all subsequent analyses were performed on male mice. We first compared the responses of interneurons and glutamatergic projection neurons by subclustering all neurons. Neuronal subtypes were identified using the Allen Brain Atlas interface “MapMyCells” to identify 4 Interneuron subclusters (*Sst, Parv, Lamp5* and *Vip* interneurons) and 8 Glutamatergic ones (L2/3, L4/5, L5, L6 intratelencephalic (IT) neurons and L5 near-projecting (NP), L5 extratelencephalic (ET), L6 corticothalamic (CT) and L6b neurons, **Fig. 1I**). To characterize the overall response of neurons to chronic hypoxia, we first performed a GSEA analysis using hallmark genes (**Fig. 1J**). Interestingly, several gene sets related to oxidative changes (i.e. *oxidative phosphorylation, hypoxia, reactive oxygen species pathways, glycolysis*), were among the top upregulated at P11. Notably, *hypoxia* gene set enrichment returned to baseline by P45, indicating a robust but transient transcriptional response. This supports the predominantly acute impact of the model and validates the use of large-scale transcriptomic analyses to assess environmental effects on cortical cell types. Of note, genes related to *oxidative phosphorylation* continued being perturbed at P45 following hypoxia, suggesting long term alteration of mitochondrial functions. Further, interferon alpha response was affected at both timepoints, although in opposite direction (decreased at P11 and increased at P45), suggesting complex effect of hypoxia on inflammation. Finally, we investigated the response of neurons subtypes to hypoxia. Mapping of DEG induced by hypoxia at P11 revealed a larger transcriptional dysregulation in pyramidal neurons than in interneurons (**Fig. 1K**). This analysis highlighted pronounced transcriptional dysregulation in L2/3 IT neurons (487 DEGs) relative to other neuronal subtypes, suggesting that late-born neurons are particularly susceptible to hypoxic conditions. This population was thus prioritized for further investigation.

### Hypoxia drives increasing mitochondrial dysregulation and fragmentation within upper cortical neurons

Given that mitochondria are highly sensitive to oxygen tension and play a key role in neuronal maturation and circuit formation^12–14^, we next examined to which extent hypoxia impacts mitochondrial biology. To this end, we performed gene set enrichment analysis (GSEA) in L2/3 neurons, at postnatal day 11 (P11) and day 45 (P45). Focusing on gene set categories related to mitochondrial processes, we noticed that the proportion of mitochondria-related gene sets increased from 10.5% (32 out of 302 gene sets) at P11 to 30% (21 out of 69 gene sets) at P45, suggesting a progressive enrichment of mitochondrial pathways among perturbed biological processes over time (**Fig. 2A**). Moreover, mitochondrial gene sets were prominently overrepresented among those commonly affected at both time points, accounting for approximately 50% (17 out of 35), indicating that sustained transcriptional alterations increasingly converge on mitochondrial function (**Fig. 2B**).

**Figure 2.**
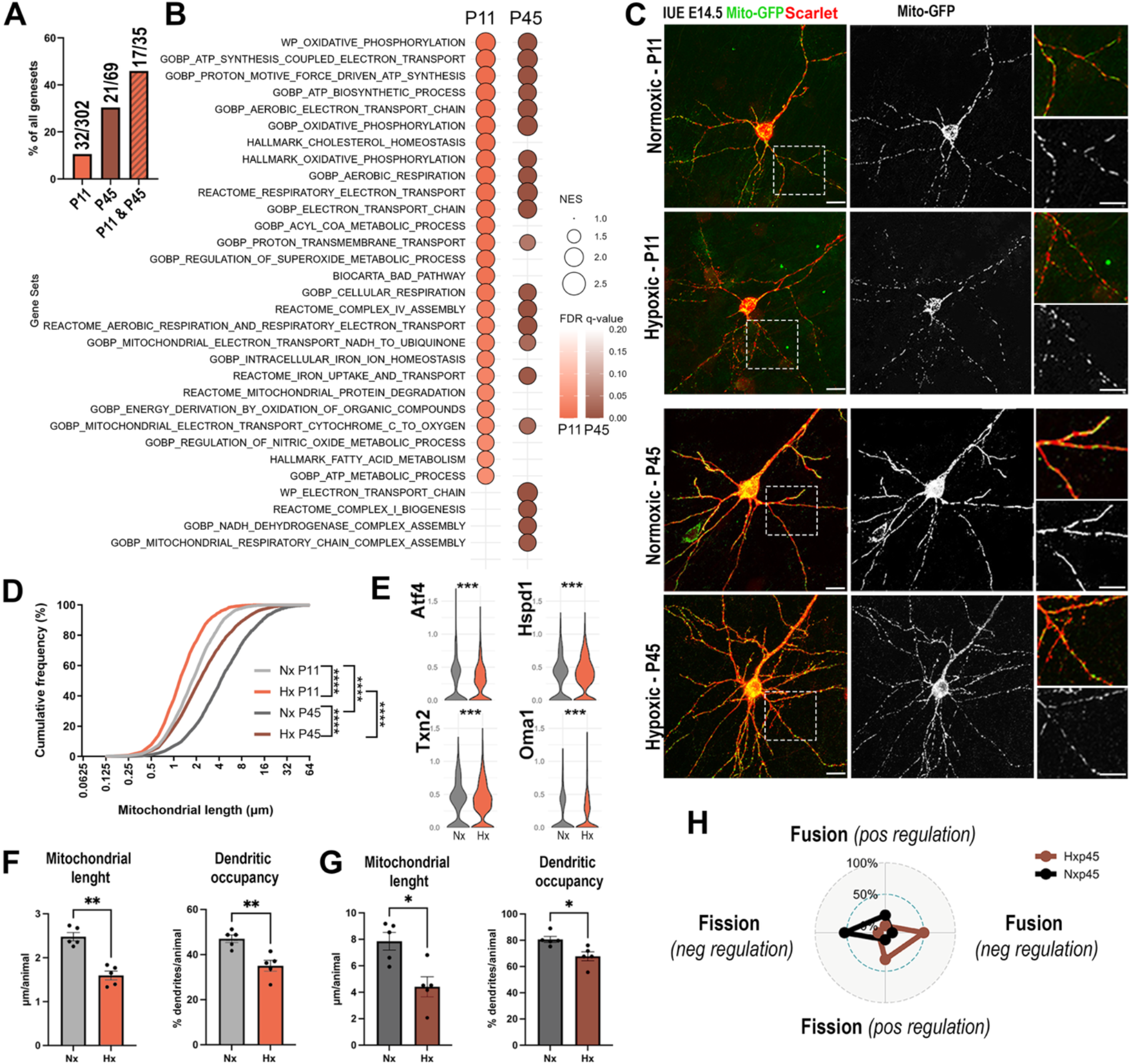
Chronic neonatal hypoxia drives long-term changes in mitochondrial gene expression accompanied by mitochondrial fragmentation. A) Proportion of mitochondria-related among all dysregulated genesets identified by GSEA analysis of layer 2/3 cortical neurons at P11 and P45 following hypoxia. B) Selected genesets reflecting pronounced and long-lasting effect of chronic hypoxia on mitochondria biology. C) Mitochondria in layer 2/3 cortical neurons dendrites were labeled by in utero electroporation of pCAG-GFP-OMM (green signal) mixed with pCAG-mScarlet-I (red signal) plasmids. Mitochondria morphology was visualized through the OMM-GFP channel within mScarlet+ electroporated neurons at P11 and P45 in normoxic and hypoxic mice. Inserts show higher magnifications of the dendritic regions indicated by boxes. Note the difference in morphology between elongated, fused mitochondria and shorter mitochondria as a function of time (P11 and P45) and hypoxia exposure. Scalebars = 10 µm. D) Cumulative distribution of mitochondria of various sizes under the two experimental conditions at P11 and P45. A shift toward shorter mitochondria is observed at the end of the hypoxic period (P11), indicating mitochondria length reduction. This phenotype persists at a longer timepoint (P45). Statistical test: ordinary one-way ANOVA with Bonferroni’s multiple comparisons test. Each quantification is from five independently electroporated pups. N_(P11Nx)_ = 263 dendrites, 2404 mitochondria, N_(P11Hx)_ = 213 dendrites, 2137 mitochondria, N_(P45Nx)_ = 313 dendrites, 2167 mitochondria, N_(P45Hx)_ = 432 dendrites, 4386 mitochondria. Number of mitochondria per dendrite ****p < 0.0001. E) Violin plots illustrating hypoxia effect on selected genes from GO terms related to activation of integrated stress signaling and mitochondrial quality control pathways. F) Quantification mitochondrial length and occupancy in dendrites of Layer 2/3 pyramidal neurons in normoxic and hypoxic mice at P11. Statistical test: unpaired t test Mann-Whitney. Data: average±SEM. N_(P11Nx)_ = 5, N_(P11Hx)_ = 5, N_(P45Nx)_ = 5, N_(P45Hx)_ = 5. G) Quantification mitochondrial length and occupancy in dendrites of Layer 2/3 pyramidal neurons in normoxic and hypoxic mice at P45. Statistical test: unpaired t test Mann-Whitney. Data: average±SEM. N_(P11Nx)_ = 5, N_(P11Hx)_ = 5, N_(P45Nx)_ = 5, N_(P45Hx)_ = 5. H) Module scores for gene sets expression involved in positive and negative regulation of mitochondria fusion and fission in Layer 2/3 pyramidal neurons at P45. *p≤ 0.05, **p≤ 0.01, ***p≤ 0.001, ****p≤ 0.0001.

To investigate the histological correlates of these transcriptional changes, we performed *in utero* electroporation of the plasmid pCAG-GFP-OMM to visualize mitochondria, along with plasmid DNA encoding cytoplasmic filler pCAG-mScarlet-I at E14.5. The pCAG-GFP-OMM plasmid allows GFP expression at the outer mitochondria membrane, thereby allowing to study mitochondria morphology and distribution along dendritic arborizations of mScarlet-I positive neurons in normoxic and hypoxic mice at both P11 and P45 (**Fig. 2C**). In dendrites, mitochondria morphology and occupancy depend on the neuronal maturation stage^25^. We could indeed confirm that at P11, mitochondria are smaller and occupy less space within proximal and distal dendritic processes of layer 2/3 PNs, when compared to P45 (**Fig. 2C, D**). Chronic neonatal hypoxia induced a reduction of mitochondria length within dendrites at P11, matched by a decrease in the percentage of occupancy (**Fig. 2D, F**). This was paralleled at the transcriptional level by a significant increase both in the proportion of neurons expressing, and in the level of transcript expression for *Atf4, Hspd1, Txn2*, and *Oma1* in hypoxic neurons (**Fig. 2E**), consistent with activation of integrated stress signaling and mitochondrial quality control pathways. Indeed, these genes are involved in *stress-responsive transcription*/*protein folding, redox homeostasis*, and *mitochondrial remodeling*, respectively, and their induction coincided with pronounced mitochondrial fragmentation.

At P45, mitochondria extended to adopt an elongated morphology and occupied more space within proximal and distal dendrites, in line with neuronal maturation^25^. In the chronic neonatal hypoxia condition, we could also observe signs of maturation and mitochondria elongation compared to the P11 condition, yet there remained a significant decrease in mitochondria length and dendrite occupancy compared to the normoxic condition (**Fig. 2D, G**). These results indicate that chronic hypoxia induces mitochondria length reduction within dendrites that persists over time, despite of a clear mitochondria maturation. To further explore mechanisms underlying persistent hypoxia-induced mitochondrial fragmentation, we performed gene ontology (GO) analysis focusing on gene sets associated with the positive and negative regulation of mitochondrial fusion and fission. Based on these gene sets, we computed module scores in P45 L2/3 IT neurons under Nx and Hx conditions. This analysis revealed strikingly distinct transcriptional profiles between Nx and Hx neurons. Thus, Nx cells displayed higher scores for gene sets associated with the positive regulation of fusion and negative regulation of fission, consistent with a less fragmented mitochondrial network. In contrast, hypoxic neurons exhibited increased scores for gene sets promoting fission and inhibiting fusion, in line with the observed persistent mitochondrial fragmentation (**Fig. 2H**).

In conclusion, we report here that chronic neonatal hypoxia shows major and persistent impact on mitochondria-related genes along with histology within layer 2/3 neurons, paralleling significant changes of their morphology within the cortex at P11 and P45.

### Chronic neonatal hypoxia perturbs L2/3 cortical neurons maturation

Given the tight coupling between mitochondrial integrity and neuronal development, we next investigated the short- and long-term impact of neonatal chronic hypoxia on the maturation of L2/3 cortical neurons. First, over representation analysis in L2/3 IT neurons at P11 revealed the central role of dysregulated genes (Log2FC = 0.25, p.adj < 0.05) in axonogenesis, as well as in synapse assembly and organization (**Fig. 3A**). Interestingly, the number of dysregulated genes remained high at P45 (i.e. 254 genes) and remained related to synapses, suggesting a long-term disorganization of neuron connectivity/communication following chronic neonatal hypoxia (**Fig. 3B**).

**Figure 3.**
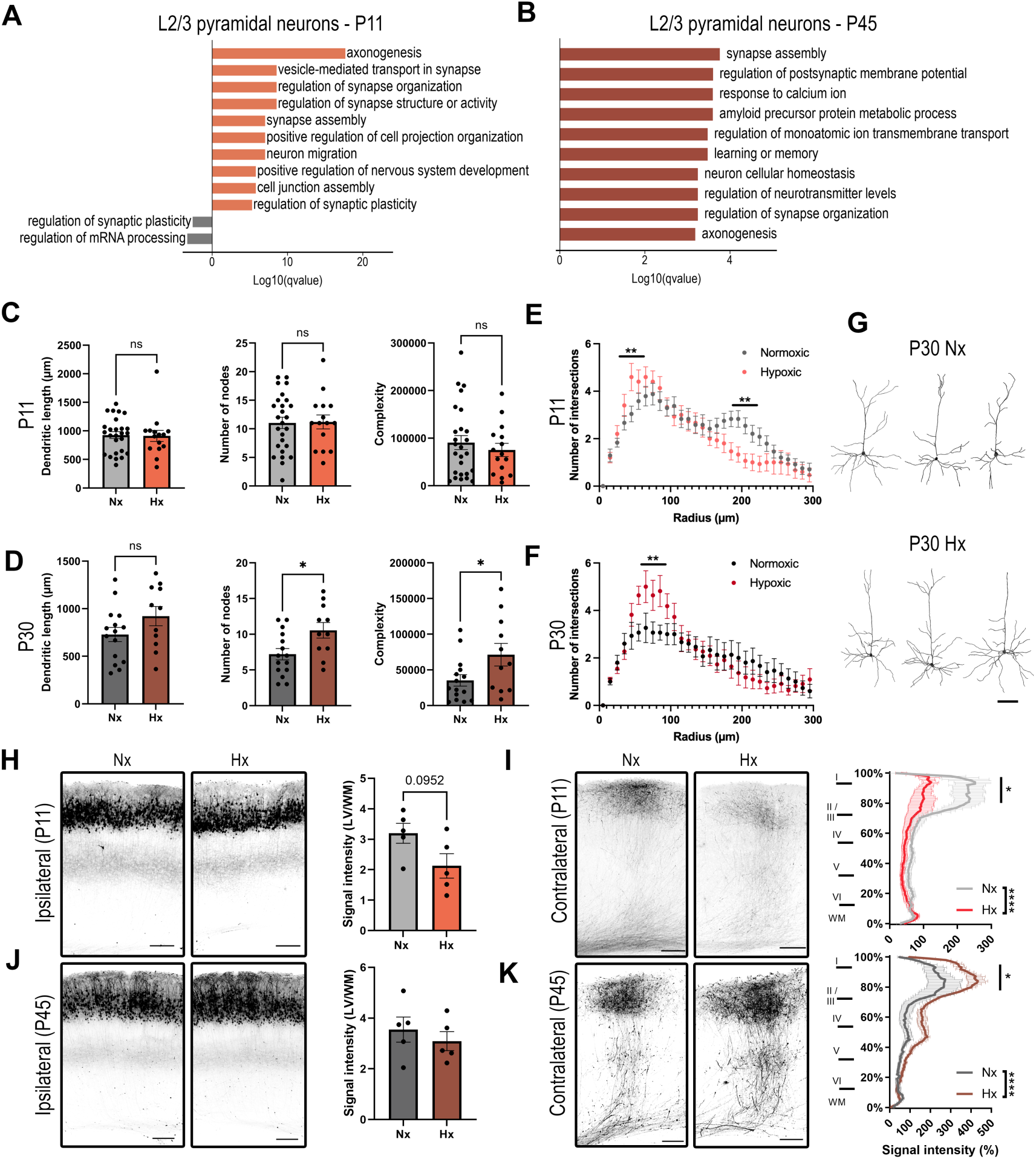
Chronic hypoxia influences Layer 2/3 neurons maturation: A-B) Gene ontology over-representation analyses on L2/3 IT neurons DEGs at P11 (A) and P45 (B). C-D) Quantifications of complexity, dendritic length and number of nodes of L2/3 neurons apical dendritic arbors at P11 (C) and following full maturation, i.e. P30 (D). E-F) Sholl analysis at P11 (E) and P30 (F) in normoxic and hypoxic conditions, highlight increase branching of apical dendrites at proximity of the cell body. Statistical test: unpaired t test Mann-Whitney (C,D); two-way ANOVA with Bonferroni’s multiple comparison (E,F). Data: average ±SEM. N_(P11Nx)_ = 24 cells, N_(P11Hx)_ = 15 cells, N_(P30Nx)_ = 15 cells, N_(P30Hx)_ = 11 cells. G) Representative 3D-reconstructions of mature normoxic and hypoxic cortical layer 2/3 pyramidal cells labeled by in utero supernova electroporation at E14.5. H-K) Representative images and corresponding quantifications of normalized pCAG-mScarlet-I fluorescence in layer 5 of the ipsilateral cortex at P11 (H) and P45 (J), as well as along the radial axis of the contralateral cortex at P11 (I) and P45 (K) between normoxic and hypoxic mice. Statistical test: unpaired t test Mann-Whitney (H,J); two-way ANOVA with Bonferroni’s multiple comparison (I,K). Data: average ±SEM. Data: average±SEM. N_(P11Nx)_ = 5, N_(P11Hx)_ = 5, N_(P45Nx)_ = 5, N_(P45Hx)_ = 5. ns not significant, *p≤ 0.05, **p≤ 0.01, ****p≤ 0.0001.

To explore the histological correlate of these transcriptional changes, we assessed the consequences of chronic hypoxia on cortical neuron maturation. We used the Supernova approach ^26^ to achieve labeling of a sparse but isochronic population of cortical neurons, to allow the optimal reconstruction of their dendritic arborization and compare them between experimental conditions. The Supernova approach relies on the *in utero* electroporation of two vectors: a CRE plasmid and a reporter plasmid (**Fig. S3A**), which ratio defines the sparsity of the recombination (**Fig. S3B-C**). We electroporated the Supernova plasmids in E14.5 mouse embryos to target the progenitors of pyramidal neurons (PNs) of superficial cortical layers 2/3 (**Fig. S3DE**). Examination of the electroporated brains from P3 on, revealed a bright expression of the reporter protein RFP, which allowed visualization of their dendritic arborization in thick coronal brain sections (**Fig. S3F**). Quantification of dendritic arbors revealed a rapid developmental increase in dendritic volume and complexity, both of which reached a plateau by P30 (**Fig. S3G**). We next assessed whether hypoxia altered dendritic maturation at P11 and P30. At P11, total dendritic length, number of nodes, and overall complexity did not differ significantly between Nx and Hx mice, suggesting that the transcriptional changes observed in L2/3 neurons were not associated with a global delay in dendritic arbor maturation (**Fig. 3C**). However, Sholl analysis on 3D pyramidal neurons reconstructions revealed early signs of dendritic remodeling (**Fig. 3E**). At this early stage, hypoxia induced a modest but significant increase in apical dendritic branching near the soma, accompanied by a reduction in branching within the apical tuft, consistent with a spatial redistribution of dendritic complexity. Dendritic arbors alterations became more pronounced over time, while they remain absent in basal dendrites (**Fig. S3H-K**). By P30, hypoxic mice exhibited a significant increase in the number of dendritic nodes and in overall arbor complexity (**Fig. 3D**), together with a stronger increase in proximal apical branching (**Fig. 3F**). Thus, rather than delaying dendritic maturation, hypoxia progressively remodeled the spatial organization and complexity of dendritic arbors (**Fig. 3G**).

Furthermore, we assessed the consequences of hypoxia on axonal projections through in *utero* electroporation of a pCAG-mScarlet-I plasmid. Electroporations were performed unilaterally, and axons of electroporated neurons were visualized based on bright expression of mScarlet, which allowed the quantitative assessment of their branching pattern within the ipsilateral and contralateral hemispheres. Analysis at P11 revealed a slight, non-significant decrease in the density of axonal branching within the ipsilateral layer 5 (**Fig. 3H**). At this early stage, axonal projections successfully crossed the corpus callosum in both normoxic and hypoxic mice to project to the contralateral cortex (**Fig. 3I**). Interestingly, a closer analysis of their branching pattern in S1/S2 cortical areas, revealed however noticeable changes following hypoxia. In particular axon branching was significantly reduced in layer 2/3 following hypoxia. Further analysis at P45 revealed a return to normoxic levels within the ipsilateral layer 5 (**Fig. 3J**). In sharp contrast, at P45 we observed a global increase in contralateral axonal branching throughout cortical layers, that reached significance in layers 2/3 in hypoxic compared to normoxic mice (**Fig. 3K**).

Together, these findings demonstrate that chronic postnatal hypoxia induces both immediate and longlasting alterations in neuronal development, characterized by an initial delay in axonal branching at P11 that subsequently translates into increased contralateral projections and enhanced dendritic and axonal complexity at later stages, ultimately indicating a profound remodeling of cortical wiring.

### Chronic neonatal hypoxia results in rewiring of cortico-cortical connectivity at adult ages

To investigate the consequence of altered L2/3 neurons maturation on cortical wiring, we used retrograde tracers to assess local and long-distance projections from L2/3 cortical neurons in hypoxic compared to normoxic mice. We performed Cholera Toxin beta-subunit (CTB) injection within the primary somatosensory cortex (S1) of P45 mice (**Fig. 4A**). At 7 days post-injection, retrogradely labeled CTB+ neurons were consistently observed in S1, secondary motor cortex (M2), secondary somatosensory cortex (S2) on the injected side. In addition, retrogradely labeled neurons were also visible in both the S1 and S2 areas, contralateral to the site of injection (CS1 and CS2). We also observed CTB+ neurons in subcortical regions corresponding exclusively to thalamic nuclei, such as the ventral-anterior lateral/medial (VAL/VM), the ventral posterior (VP) and the posterior (PO) nuclei. Together, these five brain regions (ipsilateral S1, S2, M2, contralateral S1 and S2 cortex, and ipsilateral thalamus, **Fig. 4B**) contained the vast majority of CTB+ connected neurons. Quantification of retrogradely labeled neurons throughout these brain regions in both Nx and Hx mice revealed a global increase in the total number of CTB+ neurons following chronic neonatal hypoxia that reached significance in both the ipsilateral S1 and contralateral S2 regions (**Fig. 4C**), supporting cortico-cortical hyperconnectivity. We next assessed the distribution of CTB+ neurons across cortical layers by quantifying neurons within the infragranular (layers 5/6), granular (layer 4) and supragranular layers (layers 2/3). Interestingly, hypoxia appeared to change the pattern of long-distance connectivity with the fraction of CTB+ neurons located within the infragranular layers (L5/6) of both CS1 and CS2 significantly increased in hypoxic mice, at the expense of those located within the supragranular layers (**Fig. 4D, E**).

**Figure 4.**
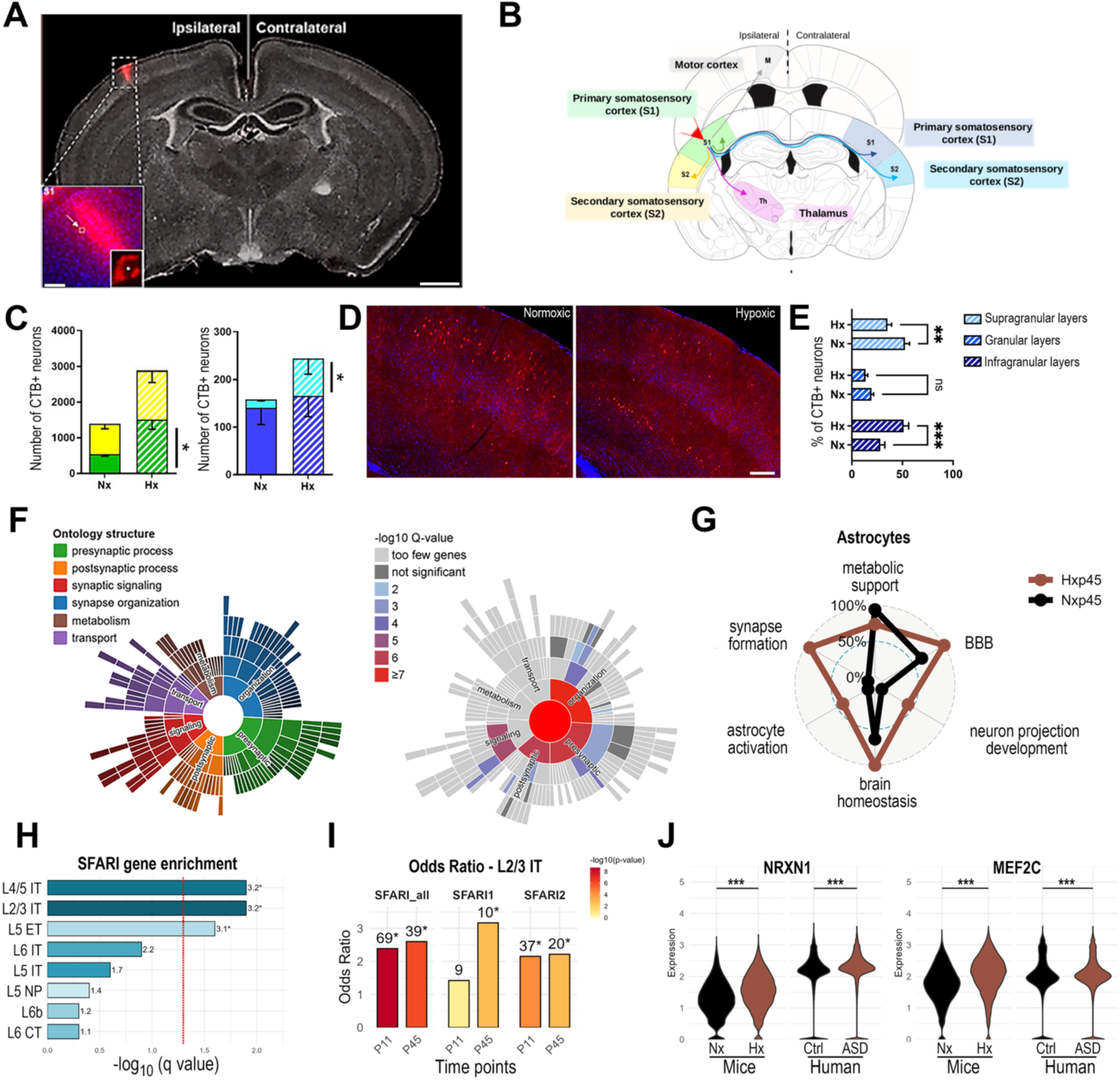
Long-term effects of chronic neonatal hypoxia on cortical connectivity. **A)** Coronal section counterstained with the nuclear marker DAPI (blue) showing the location of the site of injection of the Cholera Toxin beta-subunit (CTB) within the barrel field of the primary somatosensory cortex (S1). B) Schematic representation of the regions where CTB labeled neurons are observed: primary somatosensory cortex (S1), secondary motor cortex (M2), thalamus (Th) and secondary somatosensory cortex (S2) ipsilateral to the site of CTB injection, as well as contralateral primary and secondary somatosensory cortex (CS1, CS2). C) Quantifications of the number of CTB+ neurons in connected brain regions in normoxic vs. hypoxic P45 mice, revealing and an increased connectivity following hypoxia. Color code as described in B. D) Photomicrographs depicting CTB+ neurons in cortical connected regions, i.e. CS1, CS2 regions contralateral to the site of CTB injection. E) Quantifications within supragranular, granular and infragranular cortical layers, revealing a modified distribution of long distance connected neurons following hypoxia in CS1 merged with CS2. Statistical test in C and D: unpaired t test Mann-Whitney (C); two-way ANOVA with Bonferroni’s multiple comparison (E). Data: average±SEM. N_(P45Nx)_ = 8, N_(P45Hx)_ = 8. F) Sunburst plots for synaptic biological processes related to dysregulated genes in L2/3 neurons at P45 following hypoxia. G) Scaled module scores for gene sets expression associated with six major astrocyte functions in Nx and Hx mice at P45. H) Overlap between SFARI 1 genes and DEGs from all Glu neuron subtypes; dotted line indicates statistical significance (q < 0.05). I) Over-representation analysis of SFARI 1 and 2 genes among L2/3 IT neurons DEGs at P11 and P45. J) Illustration of 2 SFARI category 1 genes and comparison with changes observed in a human ASD dataset (Velmeshev et al., 2019) ^54^. ns not significant, *p≤ 0.05, ***p≤ 0.001. Scalebars = 1 mm (A), 150 µm (A, insert & D).

To complement these morphological analyses and further explore long-term alteration of cortical connectivity, we performed transcriptomic profiling of layer 2/3 intratelencephalic (L2/3 IT) neurons at P45, focusing on genes curated in the SynGO synapse database ^27^. Differential expression analysis revealed long-term dysregulation of genes associated with both pre- and post-synaptic components (**Fig. S4A**). Enrichment analysis highlighted affected biological processes including *“synapse organization*,” “*regulation of postsynaptic specialization*”, and “*modulation of chemical synapses*” (**Fig. 4F**). These transcriptional alterations were not accompanied by changes in dendritic spine density, either on proximal or basal dendrites (**Fig. S4B-E**). Moreover, the distribution of spine types, classified as immature (thin, filopodia-like) or mature (mushroom-shaped), remained unchanged following hypoxia (**Fig. S4F**), suggesting that the observed transcriptional shifts reflect functional rather than structural or maturational modifications. Notably, the transcriptional response to hypoxia was not restricted to neurons but also involved astrocytes. Although most astrocytic transcriptional changes were driven by age (**Fig. S4G**), a distinct subset was specifically associated with hypoxic exposure (**Fig. S4H**). Module scoring of gene sets linked to well-characterized astrocytic functions revealed a hypoxia-induced functional reprogramming, characterized by a shift from predominantly homeostatic and metabolic support programs toward pathways associated with synapse formation and neuronal projection development (**Fig. 4G**) ^28^. These changes suggest that astrocytes may actively contribute to hypoxia-induced neuronal circuits remodeling.

Finally, given the well-established association between synaptic dysfunction and neurodevelopmental disorders, we examined whether these alterations converge onto autism risk gene networks. Enrichment analysis revealed a significant over-representation of SFARI category 1 genes (**Fig. 4H**), and, to a lesser extent, category 2 genes (data not shown), specifically in intratelencephalic (IT) neurons compared to other neuronal populations. Within the IT population, upper-layer neurons, particularly those in layers 2/3 and 4/5, were more affected than deeper-layer neurons. Notably, although the overall number of DEGs declined from P11 to P45, the proportion of SFARI genes among these DEGs increased over time, suggesting that the long-term transcriptional effects of early hypoxia become progressively enriched for genes associated with autism spectrum disorder as the animals age. For instance, focusing on L2/3 IT neurons, we identified 10 SFARI category 1 and 20 SFARI category 2 genes as significantly dysregulated at P45 (**Fig. 4I**). Those included *Nrxn1* and *Mef2C*, which expression increased significantly following hypoxia at P45, similarly to results obtained in human ASD patients (**Fig. 4J**).

Together, these findings identify chronic neonatal hypoxia as a potent driver of cortical hyperconnectivity by P45, accompanied by coordinated disruption of synaptic and associated astrocytic programs and a striking enrichment of ASD-associated genes.

### Chronic postnatal hypoxia results in long-term social behavior deficits

We finally aimed at investigating the functional consequences of the delayed cortical rewiring supported by our previous observations. To this end, we first, studied the impact of chronic neonatal hypoxia on young adult (P55) and adult (P105) mice on a range of behaviors. We first performed series of behavioral assays relevant to locomotor activity- and anxiety-like behaviors. In the open field assay, we studied the spontaneous locomotor activity at P55 and P105, and found no significant differences in total distance travelled, nor in average speed between Nx and Hx mice (**Fig. S5A-C**). Next, we studied emotional reactivity, and found no significant differences in the time spent in the center zone, number of center zone entries, nor in the anxiety index, which correspond to the « Time spent in the Peripheric zone / Time spent in the center zone » (**Fig. S5D-E**). Therefore, young adult and adult hypoxic mice are active with no locomotion alterations and spend more time in the center bright lit open area of the open field, thus exhibiting no signs of anxiety-like behaviors. Further, no significant differences were detected between Nx and Hx mice in open arm duration and closed arm duration in an elevated plus maze, confirming that young adult and adult hypoxic mice did not display anxiety behaviors (**Fig. S5F-H**).

Next, we assessed the social interactions of hypoxic mice using the 3-chamber sociability assay at P55 and P105 mice. In this assay, following a habituation phase, the mouse is given a choice between a mouse and an inert object (**Fig. 5A**). As expected, Nx mice spent more time with the mouse than the object, indicating conspecific sociability. Hypoxic mice were similarly able to discriminate between a mouse and an object at P55 (**Fig. 5B**), although the time spent and number of contacts with the mouse was on average lower than in the Nx group. Furthermore, when the mice were retested at P105, this trend increased and the hypoxic mice didn’t show a preference for the mouse anymore (**Fig. 5C**). In a second test, we assessed the preference of hypoxic mice for social novelty (**Fig. S5I**). As expected, Nx mice spent significantly more time with the novel mouse. Hypoxic mice were similarly able to interact more with the novel mouse rather than the familiar one at P55, although the time spent with the novel mouse decreased when compared to mice of the Nx group (**Fig. S5J**). Furthermore, when the mice were re-tested at P105, this trend was more pronounced in the Hx group showing no statistical differences in their interactions with the familiar or novel mouse (**Fig. S5K**).

**Figure 5.**
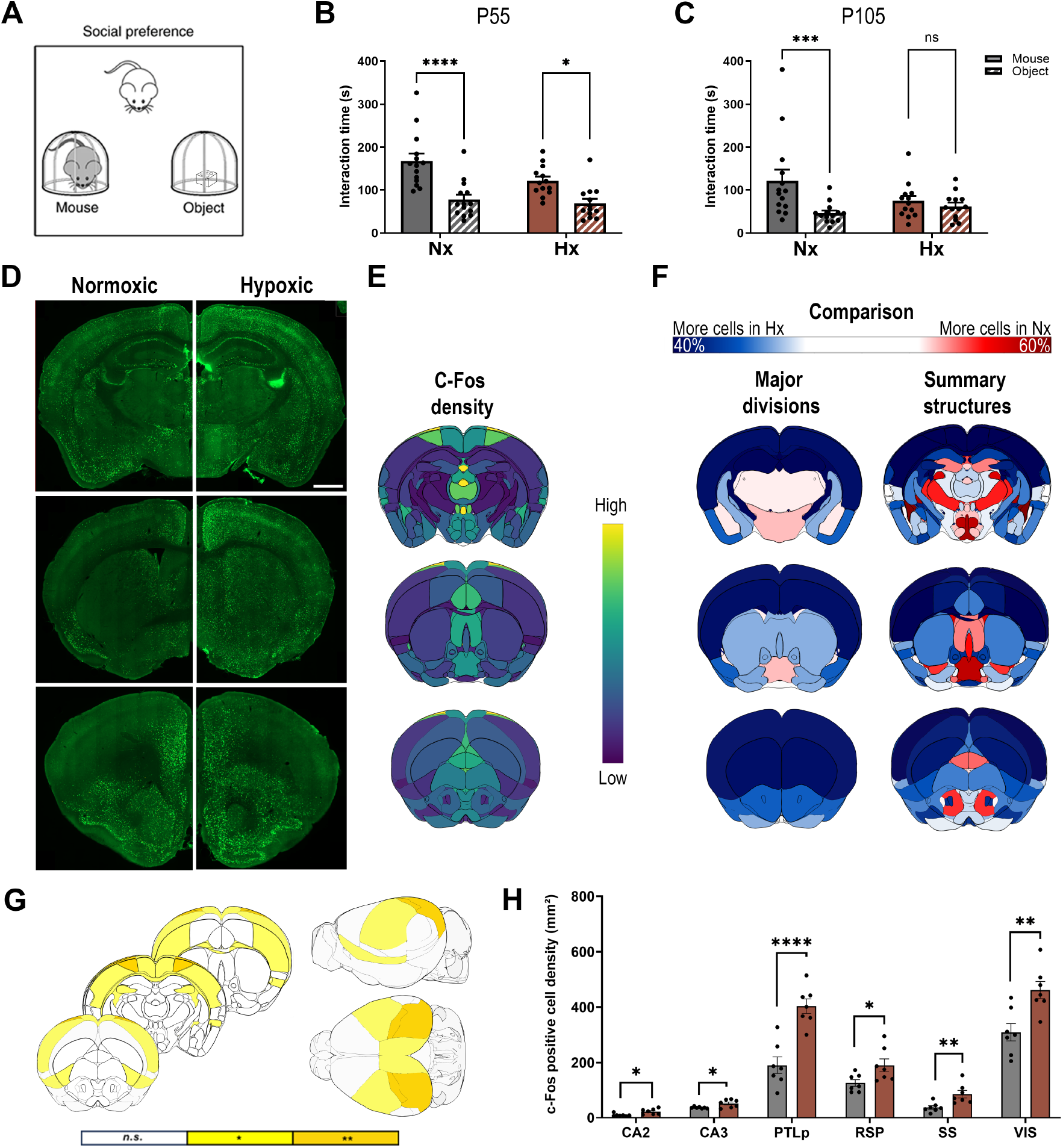
Neonatal hypoxia induces progressive social deficits associated with altered brain-wide neuronal activation. **A-C)** Sociability test schematized in A, reveals impaired social recognition at P55 (B) which amplifies at P105 (C), as demonstrated by similar interaction times with mouse and object. Statistical test: two-way ANOVA with Bonferroni’s multiple comparison. Data: average±SEM. N_(P55Nx)_ = 14, N_(P55Hx)_ = 13, N_(P105Nx)_ = 14, N_(P105Hx)_ = 13. D) Photomicrographs illustrating c-Fos immunodetection in normoxic (Nx) and hypoxic (Hx) mice at P105, 1h30 following re-exposure to a novel mouse. E) Density of c-Fos positive cells at 3 defined rostro-caudal levels of the neuraxis in normoxic mice. F) Direct comparison highlighting regions preferentially activated in Hx mice (blue gradient), when compared to Nx mice (red gradient). G) identification of brain regions showing significant differences in c-Fos proportions on a 3D model (light yellow: p<0.05; dark yellow: p<0.01). H) Selection of significant regions (n=7 mice per condition). CA2: field CA2, CA3: field CA3, VIS: visual area, PTLp: posterior Parietal Association area, RSP: retrosplenial area, SS: somatosensory area. Statistical test: two-way ANOVA with Bonferroni’s multiple comparison. Data: average±SEM. N_(Nx)_ = 7, N_(Hx)_ = 7. ns not significant, *p≤ 0.05, **p ≤ 0.01, ***p≤ 0.001, ****p≤ 0.0001

We next aimed at studying possible influences of early hypoxia onto brain activity in adulthood. To this end, we sacrificed mice 1h30 following re-exposure to a novel mouse and performed c-Fos expression mapping using serial sections encompassing the entire forebrain (**Fig. 5D**). Analyses on serial sections in normoxic mice revealed a prevalent c-Fos positive cell density within the retrosplenial cortex, the cingulate cortex, the orbital area, the prelimbic area as well as the posterior parietal associative cortex (**Fig. 5E**), consistent with the known involvement of these regions in integrating social and contextual information required for appropriate behavioral responses. Comparison of c-Fos-positive cell densities between Nx and hypoxia-exposed (Hx) mice revealed a marked imbalance in neural activity in the Hx group (**Fig. 5F**). Using a lower-resolution atlas covering major brain regions, we observed a substantial increase in c-Fos-positive cell density in the cortex, whereas I comparison subcortical structures such as the thalamus and hypothalamus exhibited a reduction (**Fig. 5F**). To further delineate the specific cortical regions affected, we utilized a more detailed atlas. This analysis revealed elevated c-Fos-positive cell densities in several cortical areas, including the posterior parietal associative cortex, retrosplenial cortex, somatosensory cortex, and visual cortex (**Fig. 5F-G**), as well as within the hippocampal formation (**Fig. 5H**). Altogether, these findings indicate heightened cortical activity in Hx mice, especially in brain regions associated with social behavior, and this altered neural activation pattern correlates with observed behavioral impairments.

## Discussion

We show that early-life oxidative stress, modelled by chronic neonatal hypoxia as observed in very preterm birth, induces both acute and persistent alterations in cortical development despite apparent macroscopic recovery. Using single-nucleus RNA sequencing, we identify widespread and long-lasting transcriptional dysregulation, with glutamatergic neurons, particularly upper-layer pyramidal neurons involved in intracortical connectivity, emerging as the most vulnerable population. These molecular changes are associated with enduring alterations in mitochondrial integrity, axonal development, and circuits wiring, and correlate with abnormal neuronal activation patterns and impaired social behaviours.

Previous studies have primarily focused on the vulnerability of GABAergic interneurons to hypoxic injury. Indeed, hypoxia has been associated with reduced density and delayed maturation of cortical interneurons, particularly parvalbumin-expressing subtypes, in both rodent models and human preterm infants^20, 29–31^. Interneuron progenitors originating from the medial ganglionic eminence are especially sensitive to hypoxia, exhibiting altered proliferation and migratory dynamics that impair cortical integration^29^. While these findings have shaped current understanding of hypoxia-induced neurodevelopmental deficits, our data indicate that glutamatergic neurons are at least equally, and in some respects more profoundly, affected than interneurons.

Notably, upper-layer glutamatergic neurons display marked vulnerability. This is consistent with recent transcriptomic work linking cortical gene expression gradients to neurodevelopmental disorders such as autism spectrum disorder and schizophrenia, where upper-layer neurons show enrichment for metabolic and synaptic gene programs associated with disease risk^32^. The selective susceptibility of these neurons may be linked to their prolonged postnatal maturation, which extends beyond the hypoxic period. It is interesting to note that a comparable developmental window of vulnerability has been described for parvalbumin interneurons, which undergo extended postnatal maturation and are particularly sensitive to environmental perturbations^33, 34^. Together, these observations support the concept that late-born neuronal populations with extended postnatal developmental trajectories are preferentially affected by chronic neonatal hypoxia.

Although prior transcriptomic studies have characterized acute effects of hypoxia-related disruptions in synaptogenesis, neurotransmission, glial maturation, and cytoskeletal organization^35^ these analyses were largely confined to the hypoxic period and performed at the whole-brain level. Consistent with previous studies, developmental “dysynchrony”, i.e. the loss of coordinated expression of genes required for synaptic maturation, has been proposed as a key feature of the premature brain^35^. Our findings extend this concept to later developmental stages, well beyond the initial insult, and further implicate additional cell types, including astrocytes. At the structural level, these molecular alterations are associated with impaired axonal development and connectivity, consistent with a miswiring phenotype. This is in agreement with diffusion tensor imaging studies in both animal models and preterm infants which have reported reduced fractional anisotropy and aberrant fiber organization following neonatal hypoxia^36–38^. Further, functional imaging in individuals with ASD supports hyper-connectivity across long- and short-range connections, that correlate to the severity of social behavior deficits ^39^, although more recent studies suggest that hyper-connectivity might be associated to a subpopulation of ASD patients ^40^. Such abnormalities likely reflect both disrupted early developmental programs and impaired postnatal refinement. Indeed, transient projections, such as callosal axons, normally undergo activity-dependent refinement during postnatal development^41, 42^, and failure of this process may contribute to the persistent connectivity defects observed here. Functionally, these alterations are associated with abnormal patterns of neuronal activation and deficits in social behaviour. The regions most affected, include the retrosplenial, cingulate, orbitofrontal, prelimbic, and posterior parietal cortices, forming a distributed network involved in contextual processing, decision-making, and social behaviour. The retrosplenial cortex supports context-dependent associative memory^43^, while the cingulate cortex integrates social and emotional information^44^. The orbitofrontal and prelimbic cortices regulate decision-making and social engagement, and the posterior parietal cortex contributes to contextual integration and behavioural adaptation ^45, 46^. Disruption across this network provides a plausible substrate for the observed behavioural deficits.

Interestingly, we show that the persistent transcriptional perturbations progressively converge on mitochondrial pathways. This suggests the presence of a lasting “mitochondrial scar,” providing a potential molecular substrate for the long-term consequences of prematurity. While acute mitochondrial dysfunction under hypoxia is well established, the long-term convergence of transcriptional changes onto mitochondrial pathways, together with sustained mitochondrial fragmentation, is more surprising. Mitochondrial dynamics, governed by the balance between fission and fusion, which balance appears to be affected on the long term by neonatal chronic hypoxia, are critical for neuronal development and survival. Excessive fission is associated with loss of mitochondrial membrane potential and neuronal death under stress conditions^47^, whereas impaired fusion leads to mitochondrial dysfunction and developmental defects^48^. Beyond energy production, mitochondria regulate calcium homeostasis, lipid metabolism, and reactive oxygen species signalling, all of which are essential for dendritic growth and synapse formation ^49–51^. Consistent with this, mitochondrial maturation has been shown to parallel neuronal differentiation and dendritic arborization ^49^, and abnormal mitochondrial morphology is a feature of several neurodevelopmental disorders, including autism spectrum disorder^52^. Our findings therefore suggest that persistent disruption of mitochondrial homeostasis represents a key mechanism linking early hypoxic injury to long-term circuit dysfunction.

Altogether, our results indicate that chronic neonatal hypoxia induces a cascade of events, from early transcriptional dysregulation to persistent mitochondrial dysfunction and circuit miswiring, that disproportionately affects upper-layer intracortical projection neurons and culminates in behavioural deficits. These findings extend current models of hypoxia-induced brain injury by highlighting the vulnerability of excitatory neurons and identifying mitochondrial dysfunction as a long-term pathological hallmark. Targeting mitochondrial homeostasis and developmental timing may therefore represent promising strategies to mitigate the neurodevelopmental consequences of prematurity.

## Material and Methods

The code for the analysis of single nuclei RNA-seq data and reproduction of all figures is available at https://github.com/OlivierRaineteauSBRI/cortex_dataset.

### Animals

Number OF1 mice (Oncins France 1, Charles River, France, MGI: 5649743) and number RjOrl:SWISS mice (Janvier labs, France) were used for the project. Mice breeding and handling were performed following the French and European legislation. All animal experiments were performed in accordance with European requirements 2010/63/UE and have been approved by the Animal Care and Use Committees of the University of Lyon CELYNE and ACCeS, under the references APAFIS #35526 and #37500 respectively.

### General design of the experiment

Mice were exposed to a hypoxic period characterized by a low O2-level concentration in the chamber (Biospherix) described in detail in ^21, 31^. Briefly, this apparatus maintains an O2 concentration in the chamber around 9.5%-10.5% using a continuous flow O2 displacement with N2. Hypoxia began at postnatal day 3 (P3) until postnatal day 11 (P11). During the hypoxic period, food and water were supplied ad libitum and mice were under 12h light-dark cycle conditions. To prevent a change of maternal behavior and to ensure a good milk quality during the hypoxic period, all mothers were replaced at P6 and P9.

The control group, named normoxic group, followed the same procedure but with a normal level of O2 in the chamber.

After hypoxia period, both groups were housed in individually ventilated cages for the rest of the experiment. At P55 and 105, all mice followed 3 behavioral tests to evaluate their locomotion, anxiety and related social behavior. One week after the last behavioral test, a simplified social test was performed 90 minutes before sacrifice to assess neuronal activity pattern by c-Fos quantifications.

### Behavorial tests

#### Locomotion: Open Field

Mice were placed in a square white Plexiglas (40 x 40 cm, homemade) open field that is divided into 16 squares (12 outer and 4 inner) of equal areas. The walls of the open field are 22 cm high. Each animal was placed in the left corner of the field and several behavioral parameters (distance travelled in the arena, average speed of mice, time spent/number of entries in center zone and anxiety index) were recorded during a 5-minute observation period and analyzed with the videotracking software Anymaze (VivoTech, Stoeling Co. Wood Dale, IL, USA).

#### Anxiety-related behaviors Elevated Plus Maze procedure

Mice were placed in an apparatus with four arms (two open without walls and two enclosed by 15 cm high walls) 40 cm long and 7 cm wide. Each animal was placed in the intersection of the four arms facing an open arm, and the time spent in each arm was recorded during a 5-minute observation period by a video-tracking system Anymaze (VivoTech, Stoeling Co. Wood Dale, IL, USA) (). Anxiety was measured as the time spent in open arms so that more time spent in open arms indicated a lower level of anxiety.

#### Three chambered social interaction tests

Social behaviors were measured by three chambers social test previously developed by Crawley’s team ^53^. Mice were placed in a three-chambered plexiglas box measuring 20cm x 40.5 cm x 22 cm. Dividing walls were clear with small semicircular openings (3.5 cm radius) allowing access into each chamber. The middle chamber was empty, and the two outer chambers contained small, round wire cages (Galaxy Cup, Spectrum Diversified Designs, Inc., Streetsboro, OH, USA) during testing. Mice were habituated to the entire apparatus without the wire cages for 10 minutes. To assess social preference, mice were returned to the middle chamber with a stranger mouse (C57BL/6J of the same sex being tested, habituated to the wire cage) in one of the wire cages in an outer compartment and a wire cage containing a red LEGO® cube in the other outer compartment. Time spent in direct contact with the novel mouse and time spent in direct contact with the novel object was recorded for 10 minutes as was number of entries in direct contact with object/mouse. For the social novelty test, mice were returned to the middle chamber, this time with the original mouse (familiar mouse) in its chamber and a new unfamiliar mouse (novel mouse) in the previously object contained chamber. Again, the number and time spent in direct contact was recorded for 10 minutes. Interval between habituation and social preference assay, and between social preference and social novelty assay was 30 seconds. The position of the object, mouse and novel mouse was assigned in a random manner.

#### C-Fos induction: Simplified three-chambered social test

Ninety minutes before sacrifice, c-Fos expression was induced by a simplified social test. In this test, each mouse followed a habituation period of 20 minutes in a cage identical to its home cage with a round wire cage similar to the previous social test. After the habituation period, a new mouse was placed in the cage for 10 minutes.

### Single nuclei isolation

Cortical tissues were minced with a razor blade to small pieces and homogenized gently using a glass douncer (Dounce tissue grinder pestle P1610, Sigma-Aldrich) by stroking 10 times (pestle A) and 5 times (pestle B) in 600 μl of cold Salty-EZ10 Lysis buffer (TrisHcl pH 7.5 10mM, NaCl 146mM, CaCl2 1mM, MgCl2 21mM, Tween20 0.03% Sigma-Aldrich P9416, BSA 0.01%, Ez buffer 10% Sigma-Aldrich NUC101, RNase inhibitor 0.2U/μl Roche, Basel, Switzerland RNAINH). After mixing gently 2-3 times, a 70 μm-strainer mesh (Pluriselect, Germany, SKU 43-10070-50) was wet with 1ml Lysis buffer in a pre-cooled 1.5 ml LoBind tube (Eppendorf® DNA LoBind tubes 0030108051, Sigma-Aldrich) and the flow was discarded through. The homogenate was then filtered and the filter was washed with 400 μl Lysis buffer. The flow was transferred to a 1.5ml LoBind tube and the nuclei were centrifuged at 500g for 5min at 4°C. The pellet was collected in 500-μl WRB2 (Tris-HCl pH 7.5 10mM, NaCl 10mM, MgCl2 3mM, BSA 1%, RNase inhibitor 0.2U/μl) and transferred in a new tube containing 500ul WRB2 for mixing/resuspension. The nuclei were centrifuged at 500g for 5min at 4°C. The supernatant was removed and a step of debris removal was added for P45 condition as follows. The pellet was resuspended in 1ml of WRB2 and the nuclei were centrifuged at 500g for 5min at 4°C. The pellet was collected in 300 μl WRB2 and passed through a 40-μm Flowmi Cell Strainer (Sigma-Aldrich, Bel-Art H13680-0040) and sort (Aria). Nuclei count was done using LUNA and encapsuled in 10x controller of 43 μl cell suspension. For the multiplexing technic, we used the 10X genomics kit.

### Bioinformatics analysis

For single-nuclei RNAseq, data were analyzed using the R software (version 4.3.2). The Seurat package (version 4.4.0) was used to filter, normalize and analyze the data. Briefly, the data were filtered by eliminating all cells with mitochondrial genome expression above 3%, and selecting nuclei in which the total number of expressed genes is inferior to 125000.

Expression levels were then log-normalized and the 3000 most variable genes were identified. Principal component analysis (PCA) and uniform manifold approximation and projection (UMAP) dimension reductions were applied to the dataset to identify cell clusters. These clusters were identified using graphbased clustering algorithm on the first 30 components and annotated according to expression of cell typespecific genes and age.

To identify enriched genes, we performed a nonparametric Wilcoxon rank sum test using the FindMarkers() function from Seurat. P value adjustment is performed using Bonferroni correction based on the total number of genes in the datasets. Genes with a P value adjusted of <0.05, at least 0.25 average fold change (log scale) were used for the overrepresentation analysis (ORA). Identification of Differentially Expressed Genes (DEGs) was based on the following criteria: min.pct = 0.1, logfc.threshold = 0.25

ORA Analysis were performed using the R package “Cluster Profiler” using the gene set databases Gene Ontology (GO) and Kyoto Encyclopedia of Genes and Genomes (KEGG). For GSEA analysis, ranked gene list were analysed using the GSEA software v4.2.3 (Broad Institute). Analyses were performed by using the following databases: hallmark gene sets (h.all.v2024.1.Hs.symbols.gmt) canonical pathways gene sets (c2.cp.v2024.1.Hs.symbols.gmt) and GO Biological Process ontology gene sets (c5.go.bp.v2024.1.Hs.symbols.gmt). Obtained results were visualized as enrichment maps in cytoscape (version 3.8.2). Clusters can be automatically defined and summarized using the AutoAnnotate Cytoscape application.

For analysis of mitochondrial fusion and fission, we calculated module scores for feature expression programs for genes of the following GO BP processes GO:0010636: *positive regulation of mitochondrial fusion*; GO:0010637: *negative regulation of mitochondrial fusion*; GO:0090141: *positive regulation of mitochondrial fission*; GO:0090258: *negative regulation of mitochondrial fission*.

To investigate synapse-related transcriptional changes, we used SynGO, an interactive, expert-curated knowledge base dedicated to synapse biology ^27^, using the Syngo-1 gene set analysis tool (https://www.syngoportal.org/syngo1/ora). All annotations in SynGO are exclusively based on peerreviewed, expert-curated literature.

For over-representation analysis of autism-associated genes, we used the “04-03-2025 release” of the SFARI Gene database (available at https://gene.sfari.org/database/human-gene/). For cross-species comparison with human data, we analyzed the dataset generated by Velmeshev et al. (2019) ^54^, available at https://cells.ucsc.edu/?ds=autism. Briefly, SFARI gene category 1 genes were selected, and their dysregulation was compared between our mouse dataset and the human dataset.

For Radar plot analysis of Astrocytes function, we performed module scores on gene sets associated with key astrocyte functions, including blood–brain barrier regulation, neuron projection development, brain homeostasis, astrocyte activation, synapse formation, and neuronal metabolic support, as previously published ^28^.

### In utero Cortical Electroporation

At E14.5, we performed in utero Cortical Electroporation as previously described ^55^. For dendritic arborizations and dendritic spines morphology quantifications, a mix containing 5ng/μl CRE plasmid “pK031” (pK031.TRE-Cre (Supernova)) and 1ug/μl reporter plasmid “pK029” (pK029.CAG-loxP-stop-loxP-RFP-ires-tTA-WPRE (Supernova)) plus 0.5% Fast Green (Dye content ≥85 %, powder F7252, Sigma-Aldrich, St. Louis, MO, USA; 1:20 ratio) was injected into one lateral ventricle. For axonal projections and mitochondria morphology study, a mix containing 1 μg/μl endotoxin-free plasmid DNA pCAG-mScarlet-I (pCAGGS-SspB-mScarlet-I), and 0.25 μg/μl endotoxin-free plasmid DNA pCAG-GFP-OMM (GFP-OMP25), plus 0.5% Fast Green (Sigma; 1:20 ratio) was injected into one lateral ventricle. Electroporation was performed using the ECM 830 electroporator (BTX) (ECM® 830 Square Wave Electroporation System, BTX Harvard Apparatus, Holliston, MA, USA) or the Electroporator NEPA21 (Nepa Gene Co., Ltd., Ichikawa, Chiba, Japan) using five pulses of 45V; 50ms length, with 950 msec interval to target cortical progenitors.

### DNA and plasmids

We used the following plasmids: The Supernova mix (CRE plasmid “pK031” (pK031.TRE-Cre (Supernova) was a gift from Takuji Iwasato (Addgene plasmid # 69136; http://n2t.net/addgene:69136; RRID:Addgene_69136) and reporter plasmid “pK029” (pK029.CAG-loxP-stoploxP-RFP-ires-tTA-WPRE (Supernova) was a gift from Takuji Iwasato (Addgene plasmid # 69138; http://n2t.net/addgene:69138; RRID:Addgene_69138), pCAG-mScarlet-I (pCAGGSSspB-mScarlet-I was a gift from Kazuhiro Aoki (Addgene plasmid # 178521; http://n2t.net/addgene:178521; RRID:Addgene_178521)), pCAG-GFP-OMM (GFPOMP25 was a gift from Gia Voeltz (Addgene plasmid # 141150; http://n2t.net/addgene:141150;RRID:Addgene_141150)). Endotoxin-free plasmid DNA was obtained using Machery Nagel midi-prep kit (NucleoBond® Midi Kit, Macherey-Nagel, Düren, Germany; Cat. No. 740420.50).

### Stereotactic injections

P45 Adult mice were anaesthetized using ¼ Xylazine (Rompun® 2%; Bayer) and ¾ Ketamine (Imalgène® 1000; Boehringer Ingelheim) mix, 1,2μl/g intraperitoneal injection, and placed in a stereotactic frame (Stoelting, Wood Dale, IL, USA). A small unilateral craniotomy (diameter: 0.5 mm) was drilled over the barrel field of the primary sensory cortex using a highspeed dental drill at the following coordinates: anteroposterior (AP) = −1.3, mediolateral (ML) = 3 mm with Bregma as reference (Paxinos et Franklin 2019). A glass pipette was heat-pulled to produce a tip of approximately 25 μm in diameter. It was then filled with Cholera Toxin beta-subunit (CTB) coupled to Alexa555 (Molecular Probes™, Thermo Fisher Scientific, Cholera Toxin Subunit B (Recombinant), Alexa Fluor™ 555 Conjugate, C-34776) and inserted in the right primary sensory cortex at the following coordinates: AP = −1.3, ML = 3, dorsoventral (DV) = −2mm with bregma as reference. 69nl of CTB was subsequently injected at a pace of 46 nl/s with a 20s interval between each injection until a total volume of 276 nl was injected using the nanoject II Auto-Nanoliter injector (Drummond™ Nanoject II Auto-Nanoliter Injector Drummond Scientific Company, Broomall, PA, USA). The glass injection pipette was left for 2 min following injection completion before being slowly removed. The wound was closed using sutures and the mouse was allowed to recover on a heating plate. Mice were left for 3 days to allow enough time for CTB to be efficiently retrogradely transported in connected neurons.

### Tissue processing

Mice electroporated with Supernova approach or injected with CTB as well as all mice, ninety minutes after the simplified social test, were euthanatized by injection of an intraperitoneal overdose of pentobarbital (EUTHASOL® vet, Virbac Animal Health) 60 mg/kg. Mice were perfused transcardially with 10 mL of a modifier Ringer solution contains 1L of Ringer, 2.5g of NaNO2 (vasodilator), 1000U Heparin (Héparine sodium salt from porcine intestinal mucosa H3393, anticoagulant; Sigma-Aldrich) and 10mg phenol red (which gives a red colored solution keeping us from mixing Ringer solution with PFA solution while perfusing mice in parallel). Mice brain were fixed with 60 mL of 4% paraformaldehyde (Paraformaldehyde 32% Aqueous Solution EM Grade, Electron Microscopy Sciences) in PBS buffer, kept on ice during the entire procedure. Brain sections were cut using vibratome (Leica VT1000 S Vibrating blade microtome, Leica Biosystems) at a frequency of 6.5 and speed of 7. For 3D neuronal reconstructions, we poduced 80μmsections with an adapted cutting plane (i.e. perpendicular to the cortical surface). This allowed capturing both the basal and apical dendritic arborization of individual neurons ^56^. For c-Fos analysis, brains were cut into sections of 50 µm of thickness. Sections were serially collected in series of 12. For the immunochemistry, 2 wells were used to ensure homogeneous sampling of the ROI and sufficient number of sections per structure according to stereological standards. For retrograde (CTB), anterograde tracing (axonal projections) and mitochondria study, we performed 50μm-sections.

Mice electroporated with the plasmid mix pCAG-mScarlet-I and pCAG-GFP-OMM were anesthetized with 5% isoflurane (VETFLURANE®, Virbac) and 2ml/min oxygen supply, and directly perfused transcardially with 2% paraformaldehyde and 0,075% Glutaraldehyde (Delta Microscopies) in PBS buffer kept on ice during the whole procedure, to preserve mitochondria integrity ^57^.

### Immunohistochemistry

#### Immunostaining for cortical layer-specific, interneuron and GFP markers

To confirm the electroporation at E14.5 of a homogeneous and isochronic Supernova+ population of cortical neurons, we performed immunodetection of specific cortical layer markers (i.e. Cux1 for superficial and Ctip2 for deep cortical layers). To study connected inhibitory neurons, we performed immunodetection of the most abundant interneuron markers in the cortex (i.e. Parvalbumin (PV) covering 55% and Vasoactive Intestinal Peptide (VIP) covering 10-15%). To quantify mitochondria within electroporated neurons, we performed immunodetection of the GFP expressed on the OMM. Free-floating sections were rinsed 3 times 5 minutes in PB 0.1M. Because Cux1 and Ctip2 markers are nuclear, we performed an antigen retrieval by incubating tissues in 0.01M citrate buffer (10mM, pH 6; Sigma-Aldrich) at 80°C for 20min under constant agitation (300 rpm). After cooling at room temperature for 20min, the sections were washed in 0.1M PB (3 x 5min). Tissues were then permeabilized, and unspecific antibody binding blocked by incubating sections for 2 hours in TNB-Tx buffer (i.e. buffer containing 0.25% Bovine Serum Albumin (BSA) (Sigma-Aldrich, A9418), 0.05% Casein (Sigma-Aldrich, C5890), 0.25% Top block (Lubio Science, SB232010), and supplemented with 0.4% triton X100 ((Sigma-Aldrich, T9284; TNB-Tx)). An incubation with primary antibodies (Ctip2: rat-Abcam#ab18465; Cux1: rabbit-Santa Cruz biotechnology#sc13024; PV: mouse-Sigma Aldrich P3088; VIP: Rabbit-Euromedex#20077; GFP: chicken-Rockland# 600-901-215) diluted in TNB-Tx buffer was then performed overnight at 4°C. Following extensive washing in PB-Tx, sections were then incubated with species-matched secondary antibodies (Donkey anti Rat Alexa Fluor 647 (Thermo Fisher Scientific, A31573); Biotinylated Donkey anti Rabbit (Jackson Immunoresearch, 711-065-152); Donkey anti Mouse Alexa Fluor 647 (Thermo Fisher Scientific, A31571); Donkey anti Rabbit Alexa Fluor 488 (Jackson Immunoresearch, 711-545-152); Goat anti-chicken Alexa Fluor 488 (Invitrogen, # A-11039) for 2 hours at room temperature. Finally, sections were counterstained with the nuclear marker 4’, 6-diamidino-2-phenylindole (DAPI, ThermoFisher Scientific, C10646, 1:5000), before being mounted on slides with Fluoromount anti-fading medium (Sigma-Aldrich). To ensure comparability between animals, care was taken to strictly perform all incubations in parallel for each experimental groups, and of the same length of time.

#### Immunostaining for c-Fos detection

Brain sections were first washed in 0.1 M PBS and 0.4% Triton X-100 ((Sigma-Aldrich, T9284; TNB-Tx) to remove the cryoprotectant. Sections were then incubated in 0.3% H2O2 for 1 hour then washed in 0.1 M PBS and 0.4% Triton X-100 (3*10 min). All sections followed an incubation in for 48 h at 4°C in 0.1 M PBS 0.4% Triton X-100 containing anti-cFos rat (c-Fos antibody, Cat# 226 017; Synaptic Systems) diluted to 1:50000 and washed in 0.1 M PBS and 0.4% Triton X-100 (3*10 min). After, sections were incubated for 2 hours in 0.1 M PBS and 0.4% Triton X-100 containing biotinylated rabbit Anti-Rat IgG antibody diluted to 1:1000 (Vector Laboratories)) and washed with 0.1 M PBS and 0.4% Triton X-100 (3*10 min). Then, sections followed by incubating for 2 hours with streptavidin (SA-HRP; Alexa Fluor™ Tyramide SuperBoost™ Kit; Life Technologies) diluted 1:1000 in 0.1 M PBS and 0.4% Triton X-100. After washing in 0.1 M PBS and 0.4% Triton X-100 (3*10 min), sections were incubated for 10 min in Alexa Fluor 488-conjugated tyramide (Molecular Probes, Eugene, OR, USA) by diluting the stock solution 1:500 in 0.0015% H2O2/amplification buffer. This reaction was terminated after 10 min by rinsing the tissue in 0.1 M PBS (3 × 10 min). To finish, sections were incubated for 5 minutes in 4',6-diamidino-2-phenylindole diluted to 1:5000 in 0.1M PB solution. Sections were then mounted onto gelatin-coated slides, dried before cover slipping.

### Images Acquisition and Analyses

#### Dendritic arborizations

In order to define optimal Supernova plasmids concentration, we quantified the rate of successful recombination among electroporated neurons on confocal image stacks acquired at 20X (N.A.: 0.7; z-step: 3μm; resolution: 1024×1024 px) on a Leica TCS SPE II confocal microscope (Leica Biosystems, RRID: SCR_002140). Recombined neurons (red) were quantified among all electroporated (green) ones, using ImageJ. For 3D reconstruction of dendritic arborization, only neurons with their cell body located in the middle of the section (i.e. 40 ± 10 μm) were selected, to acquire whole neuron’s dendritic arborizations. Image stacks covering the entire section thickness (80 μm) were acquired to include the maximum amount of dendritic arborization (20X objective, N.A: 0.7). A blind deconvolution was performed for all image stacks (Leica Las X, Autoquant™; 10 iterations). Full 3D reconstructions of the dendritic arborization were performed with the software “Neurolucida 360” (MBF Bioscience, RRID: SCR_016788). Analyses of the dendritic arborization organization were performed with companion software Neurolucida Explorer (MBF Bioscience, RRID: SCR_017348).

#### Dendritic spines

Images were acquired with laserscanning confocal microscope LSM 880 (Carl Zeiss, Germany). High resolution images of dendritic spines from Supernova electroporated neurons were acquired using the Airyscan technology (Zeiss) 100X (N.A.: 1.46; z-step: 0.1μm; resolution: 1024×1024 px). Z stacks of digital images were captured using the ZEN 2.3 software (Zeiss, RRID: SCR_013672). For the figures, the Z stacks were collapsed in one resulting picture using the maximum intensity projection function provided by the software. A deconvolution was performed for all image stacks (Huygens 20.10 (SVI); 40 iterations).

#### Mitochondria morphology

Fixed samples of 80-μm were imaged on a Nikon Ti-E microscope equipped with the C2 laser scanning confocal microscope. Nikon objectives used include 20x (0.75 N.A.) or 60x oil (1.4 N.A.). Analysis of mitochondrial length and occupancy were performed using Fiji (Image J, NIH, RRID: SCR_003070).

Axonal projections. Fixed 80μm thick sections were imaged using ZEISS Axioscan Z1 (w/Zen software; Zeiss) with objective Plan-Apochromat 20x/0.8 M27. Analyses of the axonal projection branching was performed using Fiji (Image J). To assess putative defects in axon branching, we measured RFP fluorescence intensity and normalized to RFP fluorescence intensity of axons within the corpus callosum, which is correlated to electroporation efficiency.

#### CTB+ connected neurons

Fixed samples of 50-μm were imaged using ZEISS Axioscan Z1 (w/Zen software; Zeiss, SIP 57154) with objective Plan-Apochromat 20x/0.8 M27 (5 planes of 4 microns at a depth of focus of 1.78um). Quantification of CTB+ connected neurons was performed with the software “Neuroinfo” (MBF Bioscience, RRID: 017346). Analyses of the CTB+ neurons number and distribution were performed with Neurolucida Explorer.

#### c-fos analysis

Fixed 50-µm brain sections were imaged using a ZEISS Axioscan Z1 slide scanner equipped with ZEN software (Zeiss, SIP 57154) and a Plan-Apochromat 20×/0.8 M27 objective. Five optical planes were acquired at 4-µm intervals, with a depth of focus of 1.78 µm. c-Fos-positive cells were detected using the MaxEntropy automatic thresholding method across a systematic series of sections spanning the entire forebrain (50-µm section thickness; 250-µm interval). Quantification was performed using a recently established automated workflow ^58^, which enables reproducible cell counting with minimal interindividual variability. The workflow further allows hierarchical interrogation of brain atlas annotations at different spatial scales to characterize the distribution of activated cells and compare brain-wide neuronal activation patterns between experimental groups (Nx and Hx). Significant regional differences were visualized in three dimensions using BrainRender.

### Statistical analyses

All statistical analysis and graphs were performed/created in Graphpad Prism 9 (RRID:SCR_002798). Statistical tests, p-values, and (N) numbers are presented in the figure legends. N represents number of mice unless otherwise stated. Gaussian distribution was tested using D’Agostino & Pearson’s omnibus normality test. We applied nonparametric tests when data from groups tested deviated significantly from normality. All analyses were performed on raw imaging data without any adjustments.

## Figures

Figures were assembled using Photoshop 2023 (Adobe). Images brightness and contrast were minimally adjusted. When performed, adjustments were identical for control and experimental conditions.

## Acknowledgments

We thank members of the Raineteau and Courchet laboratories for valuable comments and discussions. We are grateful to Denis Jabaudon for critical reading of the manuscript, Cyril Degletagne, Amele Ahraoui, Audrey Gibert for snRNA-Seq dataset production and help with bioinformatic analyses.

## Author contributions

Conceptualization: S.E., J.C. and O.R. Methodology: S.E., E.B., G.M.D., L.F., A.A., S.D., T.G., S.M., A.C., G.M., J.C. and O.R. Investigation: S.E., E.B., G.M.D., L.F., A.A., S.D., T.G., S.M., G.M., J.C. and O.R. Supervision: G.M, J.F.G.E., J.C. and O.R. Writing original draft: S.E., J.C. and O.R. Writing-review and editing: S.E., J.F.G.E, J.C., O.R.

## Data and materials availability

The data supporting the findings of this study are available upon reasonable request to the authors.

