## Supplementary figures for "Selective vulnerability of intracortical projection neurons drives long-term cortical circuit rewiring after perinatal hypoxia"

This document contains Supplementary Figures 1 to 5, which provide additional support for the main findings.

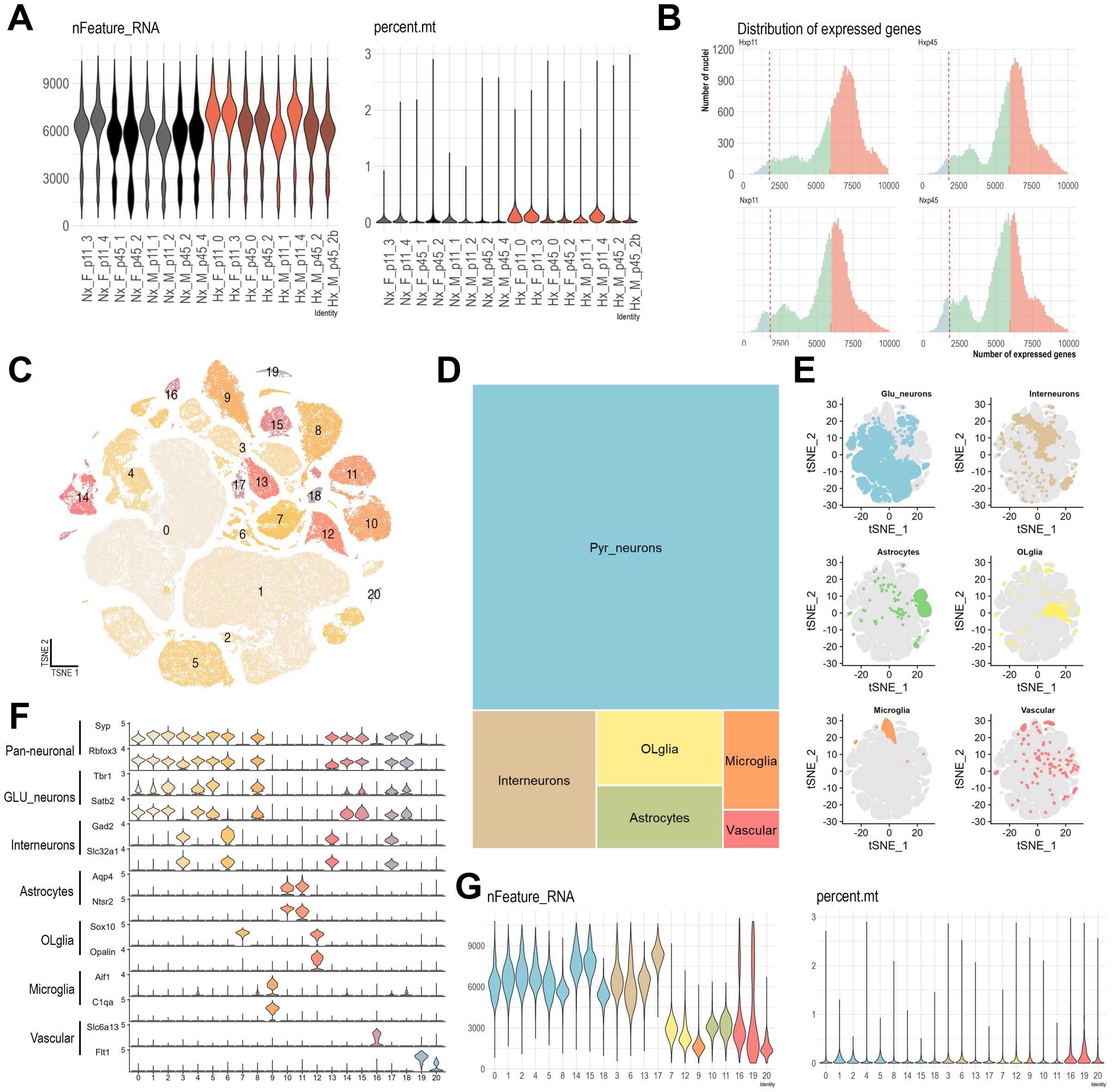


<u>***Suppl Fig. 1***</u>*, related to figure 1*<u>***:***</u> <u>***QC of single nuclei datasets***</u>*.* ***A)*** *quality control all samples, split by conditions.* ***B)*** *distribution of expressed genes between timepoints and conditions.* ***C)*** *Tsne showing all cells and clusters at dims 1:30 and low resolution (0.1) revealing 21 clusters of different sizes.* ***D)*** *Treemap representing the proportion of the major cell types present in our dataset.* ***E)*** *Combined expression of key markers on t-SNEs (one combination of markers per major cell type).* ***F)*** *Violin plots showing expression of key markers used in E and relevance to identified cell types. G) Number of detected genes and mitochondrial genes in all samples, split by identified cell types.*

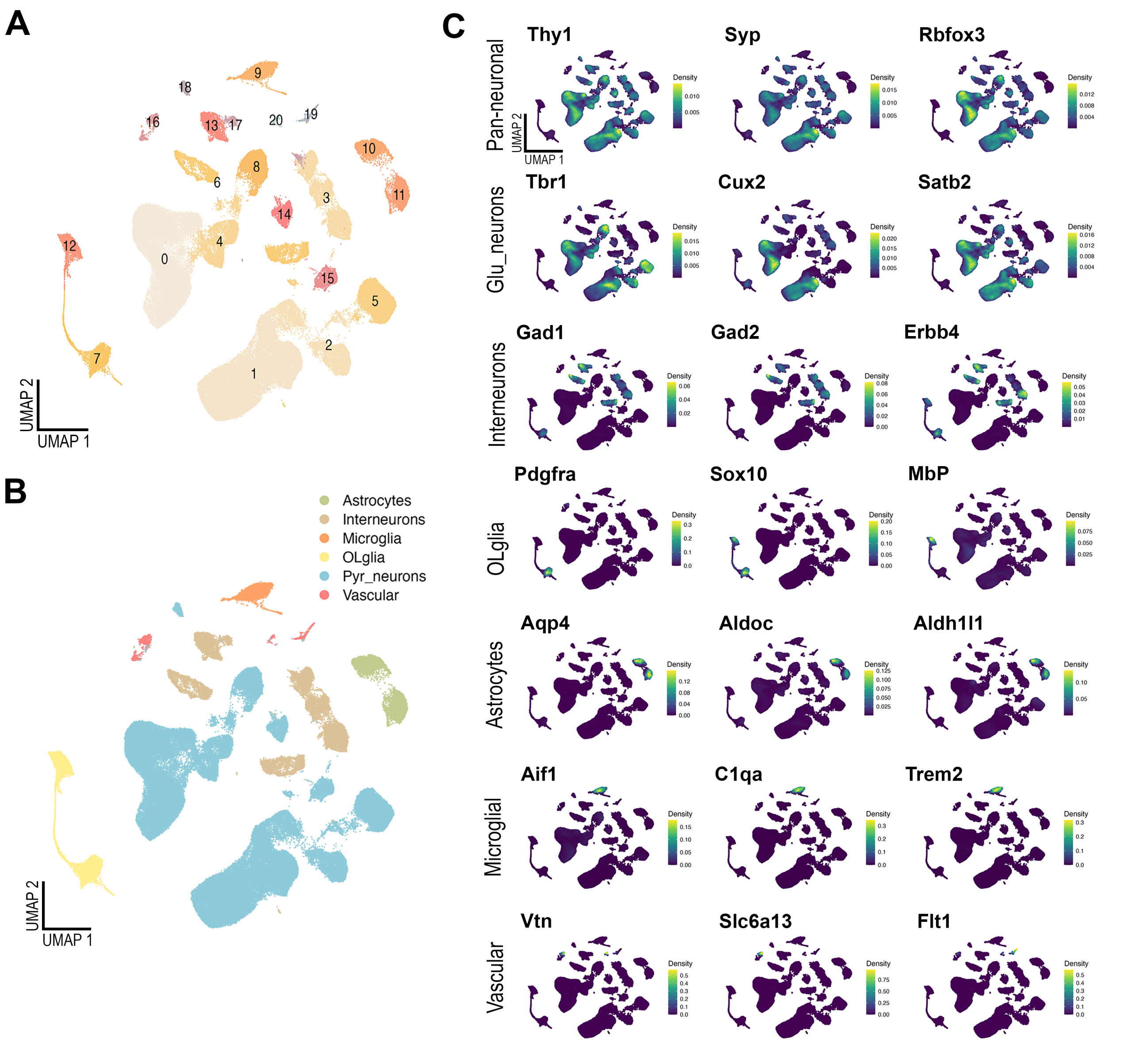


__Suppl Fig. 2__, *related to figure 1*__:__ ***Markers defining cell types of the postnatal cortex. A)*** *Clustering of all cells following QC. Umap plot produced at dims 1:30 with low resolution (0.1) revealing 21 clusters of different sizes* ***B-C)*** *Cluster identity was defined by select marker expression (C) as well as by querying the mousebrain.org adolescence atlas based on Top10 enriched genes/cluster.*

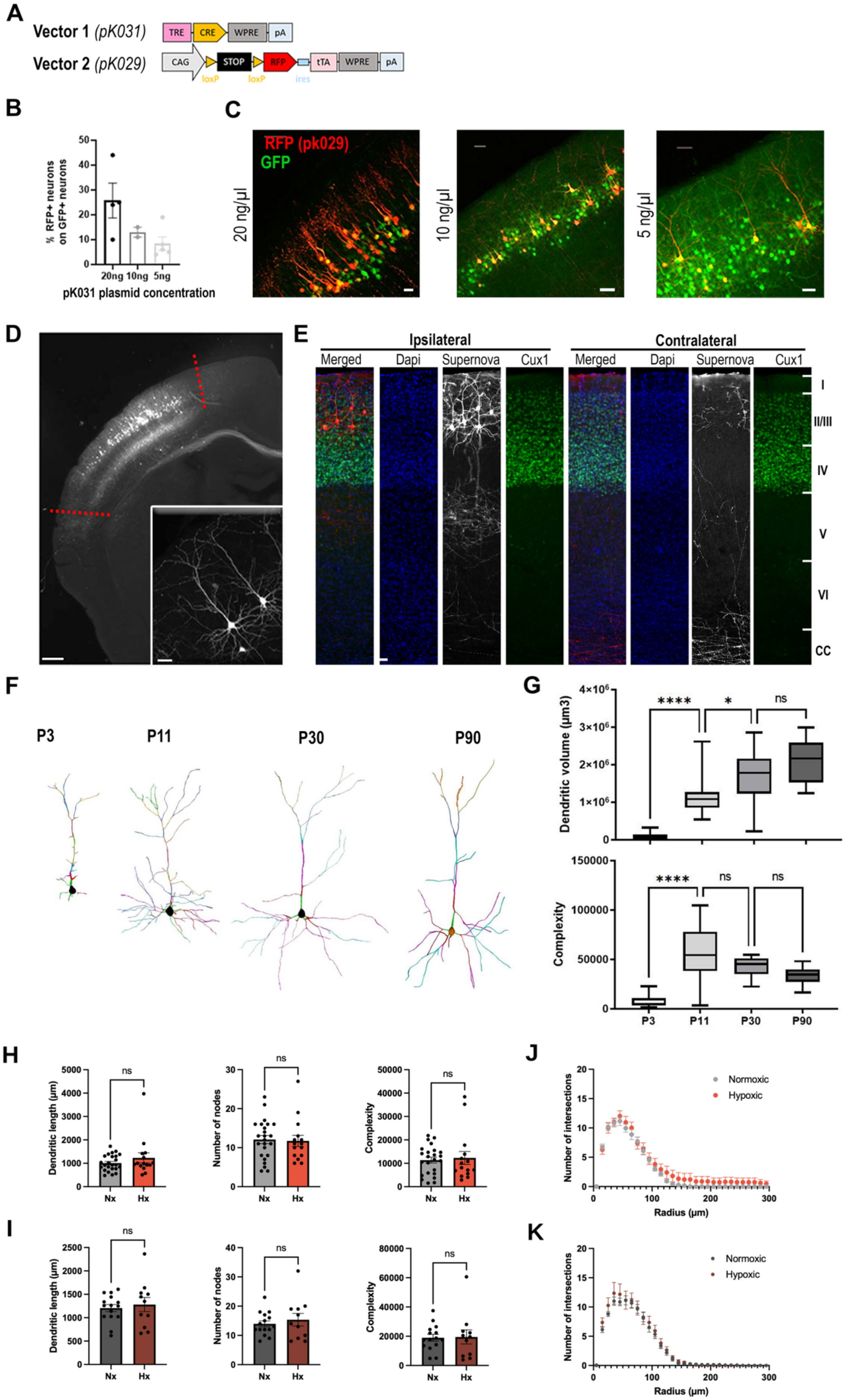
A
Vector 1 (pK031)
TRE
CRE
WPRE
pA
Vector 2 (pK029)
CAG
STOP
RFP
tTA
WPRE
pA
loxP
loxP
ires
B
% RFP+ neurons on GFP+ neurons
20ng 10ng 5ng
pK031 plasmid concentration
C
RFP (pk029)
GFP
20 ng/µl
10 ng/µl
5 ng/µl
D
E
Ipsilateral
Contralateral
Merged
Dapi
Supernova
Cux1
I
II/III
IV
V
VI
CC
F
P3
P11
P30
P90
G
Dendritic volume (µm3)
****
*
ns
Complexity
****
ns
ns
P3
P11
P30
P90
H
Dendritic length (µm)
Number of nodes
Complexity
ns
Nx
Hx
J
Number of intersections
Normoxic
Hypoxic
Radius (µm)
I
Dendritic length (µm)
Number of nodes
Complexity
ns
Nx
Hx
K
Number of intersections
Normoxic
Hypoxic
Radius (µm)

*__Suppl Fig. 3__, related to figure 3*__:__ ***Establishment of Supernova approach for sparse labeling and analysis of dendritic maturation of upper-layer pyramidal neurons A)*** *The Supernova approach is based on the use of 2 vectors: i.e. the CRE plasmid pk031 and the reporter plasmid pk029. Abbreviations: TRE: tetracycline response element; CRE: CRE recombinase; WRPE: Woodchuck Hepatitis Virus (WHP) Posttranscriptional Regulatory Element; pA: polyadenylation; CAG: CMV early enhancer/chicken β actin promoter; LoxP: site-specific recombinase; ires: internal ribosomal entry site; RFP: fluorescent protein; tTA: tetracycline transactivator. Initially, the leakage of the TRE promoter drives weak but above threshold CRE expression in a sparse population of transfected neurons that is sufficient for excision of the loxP sites and therefore RFP and tTA expression in a limited number of cells. Within these cells, tTA then binds TRE to promote expression of CRE, thereby creating a positive amplification loop allowing high and persisting expression of the reporter gene RFP.* ***B-C)*** *Representative quantifications (B) and immunostainings (C) of the proportion of recombined (RFP+/GFP+) among all electroporated cells (GFP+) when varying the concentration of the pK031 plasmid, allowing scaling of the approach for sparse neuron labeling. Data: average±SEM.* $N_{(20ng)}=4$, $N_{(10ng)}=2$, $N_{(5ng)}=5$. ***D)*** *Confocal image at P30, showing that this approach allows to visualize sparsely labeled neurons within the cortex as well as the full extent of dendritic arborization in exquisite details, up to dendritic spines. Scale bars: 250-µm and 20-µm (insert).* ***E)*** *Immunostaining showing localization of supernova+ neurons within the upper, Cux1+ layers of the cortex. Cortical layers are indicated I to VI, as well as corpus callosum (CC)* ***F)*** *Representative 3D (Neurolucida 360–based) reconstructions of individual pyramidal neurons at P3, P11, P30, and P90.* ***G)*** *Quantifications revealing a progressive increase in dendritic volume and complexity. Notably, dendritic arbors reach a mature and stable state by P30, with no further significant changes observed at later time points, supporting the use of this stage for subsequent analyses. Statistical test: One-way ANOVA with Bonferroni's multiple comparison. Data: average±SEM.* $N_{(P3)}=29$ *cells,* $N_{(P11)}=20$ *cells,* $N_{(P30)}=14$ *cells,* $N_{(P90)}=6$ *cells. ns not significant, *p≤0.05, ****p≤0.0001.* ***H-I)*** *Quantifications of complexity, dendritic length and number of nodes of basal L2/3 neurons dendritic arbors at P11 (H) and following full maturation, i.e. P30 (I)*. ***J-K)*** *Sholl analysis at P11 (J) and P30 (K) of basal dendrites in normoxic and hypoxic conditions. Statistical test: unpaired t test Mann-Whitney (H,I); two-way ANOVA with Bonferroni's multiple comparison (J,K). Data: average±SEM.* $N_{(P11Nx)}=24$ *cells,* $N_{(P11Hx)}=15$ *cells,* $N_{(P30Nx)}=15$ *cells,* $N_{(P30Hx)}=11$ *cells.*

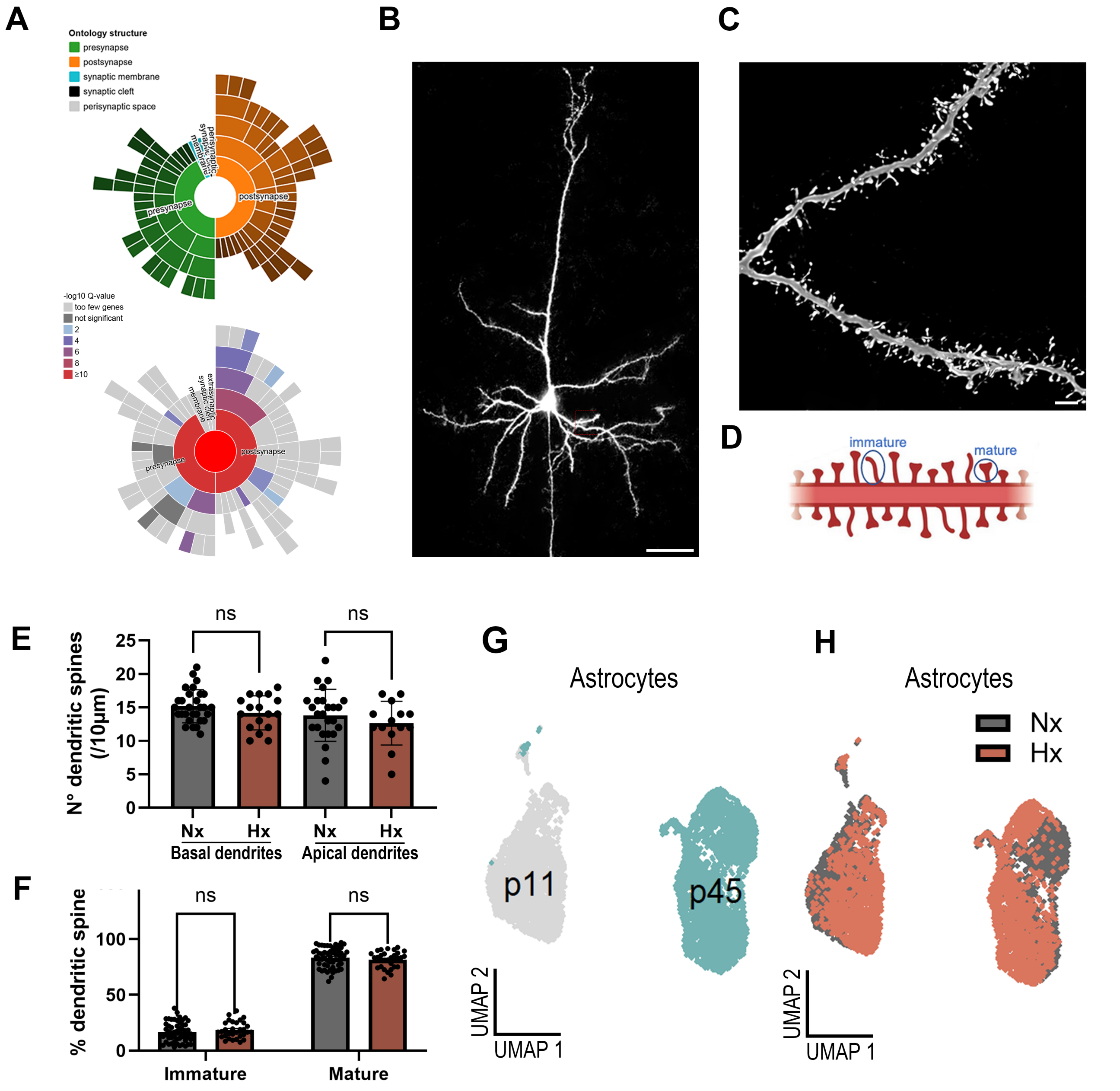


*__Suppl Fig 4__, related to figure 3 & 4: **Long-term effects of chronic neonatal hypoxia on dendritic spines and astrocytic transcriptome. A-C)** Photomicrographs of supernova+ neuron (A) and dendritic arbors (B), illustrating the identification of dendritic spines and their classifiation in mature and immature spines (C). Scale bars: 50 µm in A and 3 µm in B. **D-E)** Quantification of dendritic spines density (D) and immature/mature morphologies (E) at P30, showing no differences between experimental conditions. Statistical test: Two-way ANOVA with Bonferroni's multiple comparison. Data: average±SEM. $N_{(Nx\ basal)}$=29 cells, $N_{(Hx\ basal)}$=17 cells, $N_{(Nx\ apical)}$=25 cells, $N_{(Hx\ apical)}$=14 cells. **F)** Sunburst plots for synaptic location related to dysregulated genes in L2/3 neurons at P45 following hypoxia. **G-H)** UMAP plots of subclustered astrocytes identified by age or condition. ns not significant.*

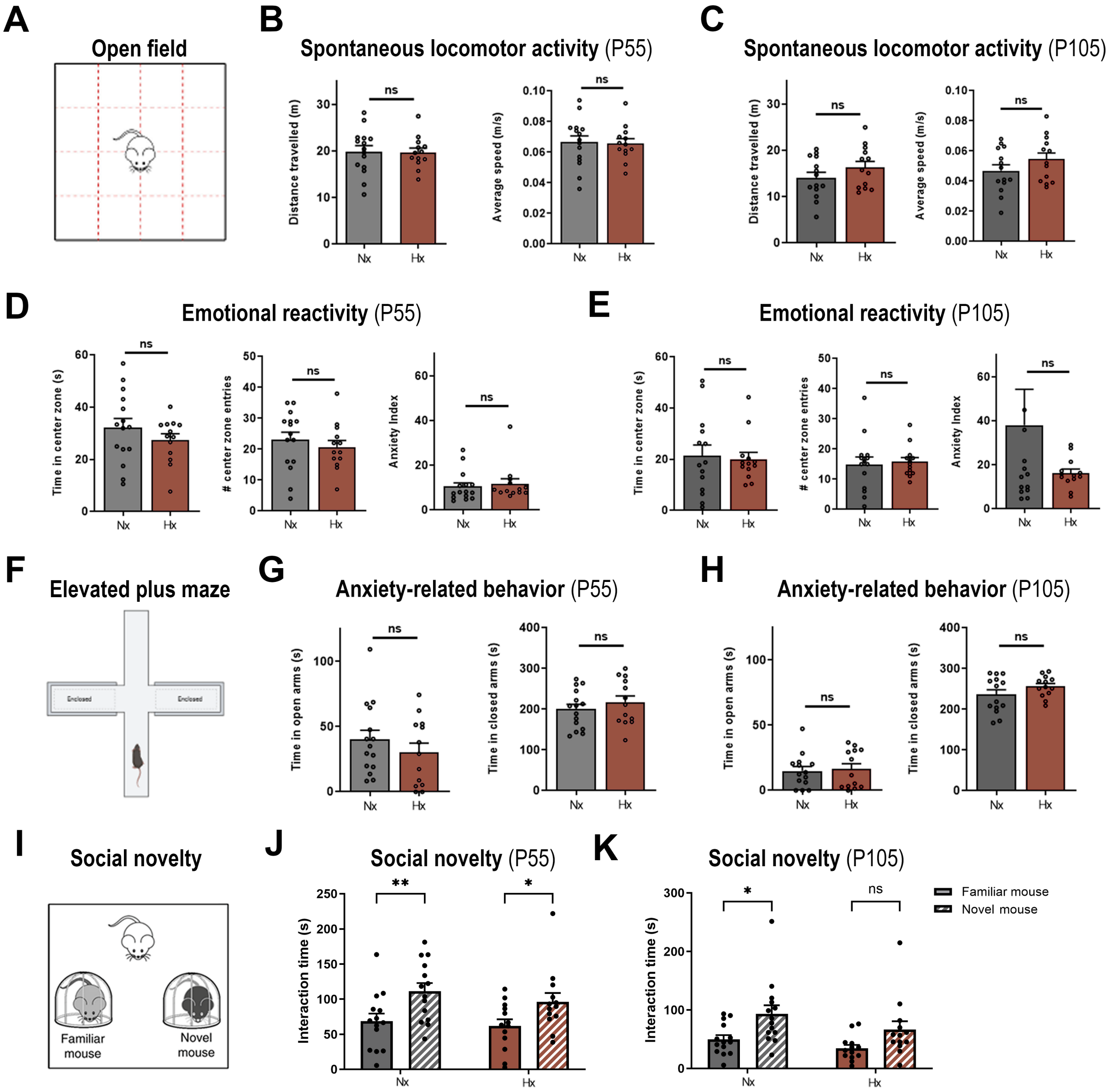


<u>*Suppl Fig. 5*</u>*, related to figure 5*: ***Neonatal hypoxia induces selective social deficits in adulthood. A-E)*** *Open filed behavioral tests (schematized in A) were performed to evaluate locomotion activity and anxiety at 55 days of age (P55, B) and were repeated at 105 days of age (P105, C). For the locomotion, distance travelled in meter were measured in mice exposed to normoxia (normoxic) and hypoxia environment (hypoxic). For the anxiety level, time spent in second in the open arms of an elevated plus maze was measured at P55 (D) and P105 (E).* ***F-H)*** *Elevated plus maze behavioral tests (schematized in F) were performed to evaluate anxiety at 55 days of age (P55, F) and were repeated at 105 days of age (P105, H), as quantified by the time spend in open vs closed arms.* ***I-K)*** *Social novelty was tested in a three-chamber test (schematized in I) at 55 days of age (P55, J) and were repeated at 105 days of age (P105, K), as quantified by the time spend to explore familiar vs novel mice. Statistical test: two-way ANOVA with Bonferroni's multiple comparison. Data: average±SEM.* $N_{(P55Nx)}=14$, $N_{(P55Hx)}=13$, $N_{(P105Nx)}=14$, $N_{(P105Hx)}=13$. *ns not significant.*
